# Mitochondrial networks of microglia adapt in a sex-specific manner upon injury-induced stress and UCP2 knockout

**DOI:** 10.1101/2022.08.22.504738

**Authors:** Margaret E Maes, Gloria Colombo, Florianne E. Schoot Uiterkamp, Felix Sternberg, Alessandro Venturino, Elena E Pohl, Sandra Siegert

**Affiliations:** Institute of Science and Technology Austria (ISTA), Am Campus 1, 3400 Klosterneuburg, Austria; Current: Department of Biomedical Sciences, University of Lausanne (Unil), Rue du Bugnon 7, 1005 Lausanne, Switzerland; Institute of Physiology, Pathophysiology and Biophysics, University of Veterinary Medicine, Veterinärplatz 1, 1210 Vienna, Austria

**Keywords:** Microglia, mitochondria, *in vitro* vs *in vivo*, UCP2, stress, hyperfusion, sex, optic nerve crush

## Abstract

Balanced and dynamic mitochondrial networks are essential for cell survival and function. Mitochondrial networks remodel their connectivity, content and subcellular localization to support optimized energy production under conditions of increased stress. *In vivo*, stressors can arise from the environment, such as in neuronal injury, or from mutation-induced cellular dysfunction. Cells programmed to identify and respond to these stress signals, like microglia, rely on optimized mitochondrial function, however we know very little about mitochondrial networks of microglia *in vivo* or their adaptation to environmental or cellular stressors.

Here, we define the mitochondrial networks of retinal microglia in physiological conditions *in vivo* and evaluate network alterations by taking advantage of a microglia-selective mitochondria-labeled mouse model. First, we demonstrate significant differences in the mitochondrial networks of microglia *in vivo* and *in vitro.* Then, we induced neuronal injury in the *in vivo* environment using optic nerve crush, where responsive microglia exhibit more fragmented mitochondrial networks with increased content and perinuclear localization, supporting a state of increased cellular stress. Surprisingly, when we selectively increase cellular stress by knocking out the mitochondria-associated gene uncoupling protein 2 (UCP2), only male UCP2^KO^ microglia establish a hyperfused mitochondrial network after injury, indicating sex differences in microglial stress mitigation. Ovariectomy in UCP2^KO^ females elicits a shift toward the male hyperfused mitochondrial phenotype suggesting that circulating estrogens are a contributing factor to the differences in microglial stress mitigation.

## Introduction

Mitochondria have been classically defined as autonomous organelles that support eukaryotic cell function by providing energy through cellular respiration (Giacomello et al., 2020). In recent years, this traditional view has expanded to include several mechanisms of mitochondrial-mediated cellular signaling such as, inter-organelle contacts, mitochondrial-derived vesicle trafficking and release of metabolites or mitochondrial DNA (Collier et al., 2023; Kornmann and Walter, 2010; Sugiura et al., 2014; West and Shadel, 2017). In line with these observations, changes in cellular metabolism and mitochondrial-associated gene signatures have been frequently associated to pathological conditions, highlighting mitochondria as key players in cell signaling and disease (Collier et al., 2023).

In physiological conditions, mitochondrial networks constantly remodel through the process of organelle fission and fusion, termed mitochondrial dynamics (Giacomello et al., 2020). Mitochondrial fusion supports matrix content exchange and/or enhanced ATP production, while fission assists redistribution and elimination of dysfunctional organelles (Youle and van der Bliek, 2012). Cellular stress, such as increased reactive oxygen species (ROS), leads to imbalanced mitochondrial dynamics often resulting in a more fragmented network. This is reflected in an increased number of small-volume, spherical mitochondria (Eisner et al., 2018) and expedites removal of damaged mitochondria in a process termed mitophagy. To compensate for mitophagy, mitochondrial biogenesis occurs which induces protein synthesis allowing the organelles to increase in size, leading to transient alterations in cellular mitochondrial content (Palikaras et al., 2015). Fragmented mitochondrial networks also facilitate organelle trafficking and redistribution to different subcellular compartments, which can influence cytoplasmic and nuclear calcium loads and transcriptional activity (Al-Mehdi et al., 2012; Kasahara et al., 2013; Park et al., 2001). Under certain stress conditions, mitochondria can also form highly connected, hyperfused networks, which increase membrane potential and ATP production helping mitigate stress and promotes cell survival (Gomes et al., 2011; Rambold et al., 2011; Tondera et al., 2009). Therefore, the adaptations of mitochondrial network connectivity, content, and localization provide valuable insight into cellular stress and energy demands.

Degenerative disease environments increase cellular stress, and failure to mitigate this stress results in its accumulation and cellular damage (Simonian and Coyle, 1996). This is particularly relevant in the context of immune cells, whose primary role is to detect and rapidly react to perturbations in their environment (Caputa et al., 2019; Collier et al., 2023; Rambold and Pearce, 2018). Microglia, the resident immune cells of the brain, respond to environmental cues through morphological, transcriptional, and metabolic adaptations (Baik et al., 2019; Hanisch and Kettenmann, 2007; Salter and Stevens, 2017). Several studies have shown that mitochondria are essential to microglial surveillance and response. Mitochondrial network fragmentation is required for the metabolic shift to glycolysis in responsive microglia *in vitro* (Nair et al., 2019). This was corroborated in cell isolation studies where aberrant metabolic reprogramming was identified as the cause for microglial dysfunction in Alzheimer’s mouse models (Baik et al., 2019). Microglia are able to adapt to available energy sources to maintain their surveillance functions (Bernier et al., 2020), however in germ-free mice where fuel sources are scarce, their mitochondrial function diminishes which is reflected in increased mitochondrial mass, ROS and reduced numbers (Erny et al., 2021).

Therefore, mitochondrial network adaptations are necessary for effective microglial responses, and provide important insights into cellular stress and metabolic changes that could predict disease-related phenotypes in the neuronal environment.

However, nearly all of the evidence investigating mitochondrial adaptations are from *in vitro* or cell isolation studies (Baik et al., 2019; Erny et al., 2021; Nair et al., 2019; Park et al., 2013; Scheiblich et al., 2021). Microglia isolated from their environment quickly transition to a responsive phenotype, preventing us from deciphering whether novel insights result from the microglia within the tissue environment or from artifacts arising due to the additional stress of tissue dissociation, processing, or the primary culture environment (Bakina et al., 2023; Gosselin et al., 2017; Marsh et al., 2022). Furthermore, spatial information and environmental factors like sex are lost in cell isolation studies, which are important contributing factors known to influence microglia (Colombo et al., 2022). A major limitation in addressing mitochondria in microglia *in vivo* is the lack of tools to effectively visualize mitochondrial networks specifically in microglia.

Here, we generated a mouse model that selectively labels mitochondria in microglia. First, we established the differences between *in vitro* and *in vivo* mitochondrial networks. Then, we analyzed spatially-isolated microglial populations in the brain using the retina as a model. Here, microglia predominately reside in inner– and outer plexiform layers (IPL and OPL, respectively), and respond to optic nerve crush (ONC) which induces axonal injury of retinal neurons distant from the retinal tissue (Bodeutsch et al., 1999; Lehmann et al., 2010; Li et al., 1999). We show that after ONC, microglia have more fragmented mitochondrial networks with increased content per cell and altered localization in IPL microglia, while OPL microglia further from the dying neurons only minimally respond. Next, we manipulated cellular stress in microglia by selectively depleting uncoupling protein 2 (UCP2), a mitochondria-localized gene associated with stress mitigation and microglial function (Ježek et al., 2018; Kim et al., 2019). UCP2-depleted IPL microglia responded similarly to wildtype after ONC with the exception of mitochondrial number. We uncovered a sex-specific difference in stress mitigation, where male UCP2^KO^ microglia exhibited hyperfused mitochondrial networks after ONC. Ovariectomized UCP2^KO^ females represented a similar mitochondrial phenotype suggesting that circulating estrogens influence this effect, and highlighting that there are sex-based differences in microglial stress mitigation.

## Results

### Mitochondrial networks of microglia differ between *in vitro* and *in vivo* environments

Visualizing mitochondrial networks of retinal microglia by immunostaining for markers such as TOMM20 (translocase of outer mitochondria membrane 20) results in a dense and ill-defined labeling in the OPL and IPL, where microglia predominantly reside (**Figure 1A**). Thus, we took advantage of the PhAM^fl/fl^ mouse, which carries a floxed transgene encoding a mitochondrial-localized Dendra2 fluorophore (Pham et al., 2012). We crossed this mouse with the *Cx3cr1*^CreERT2^ mouse expressing tamoxifen-inducible Cre recombinase in cells of myeloid origin (Yona et al., 2013). After tamoxifen administration, *Cx3cr1*^CreERT2/+^/PhAM^fl/fl^ mice, hereafter referred to as wildtype (WT) mice (**Figure 1B**), microglia labeled with IBA1 (ionized calcium binding adaptor molecule 1) showed selective expression of the mito-Dendra2 fluorophore (**Figure 1C**). We verified mito-Dendra2 localization at the mitochondria of microglia by co-labeling with TOMM20 in retinal sections (**Figure 1D**), as well as in 4-hydroxytamoxifen treated primary mixed glial cultures prepared from WT pups (**Figure 1E**).

**Figure 1.**
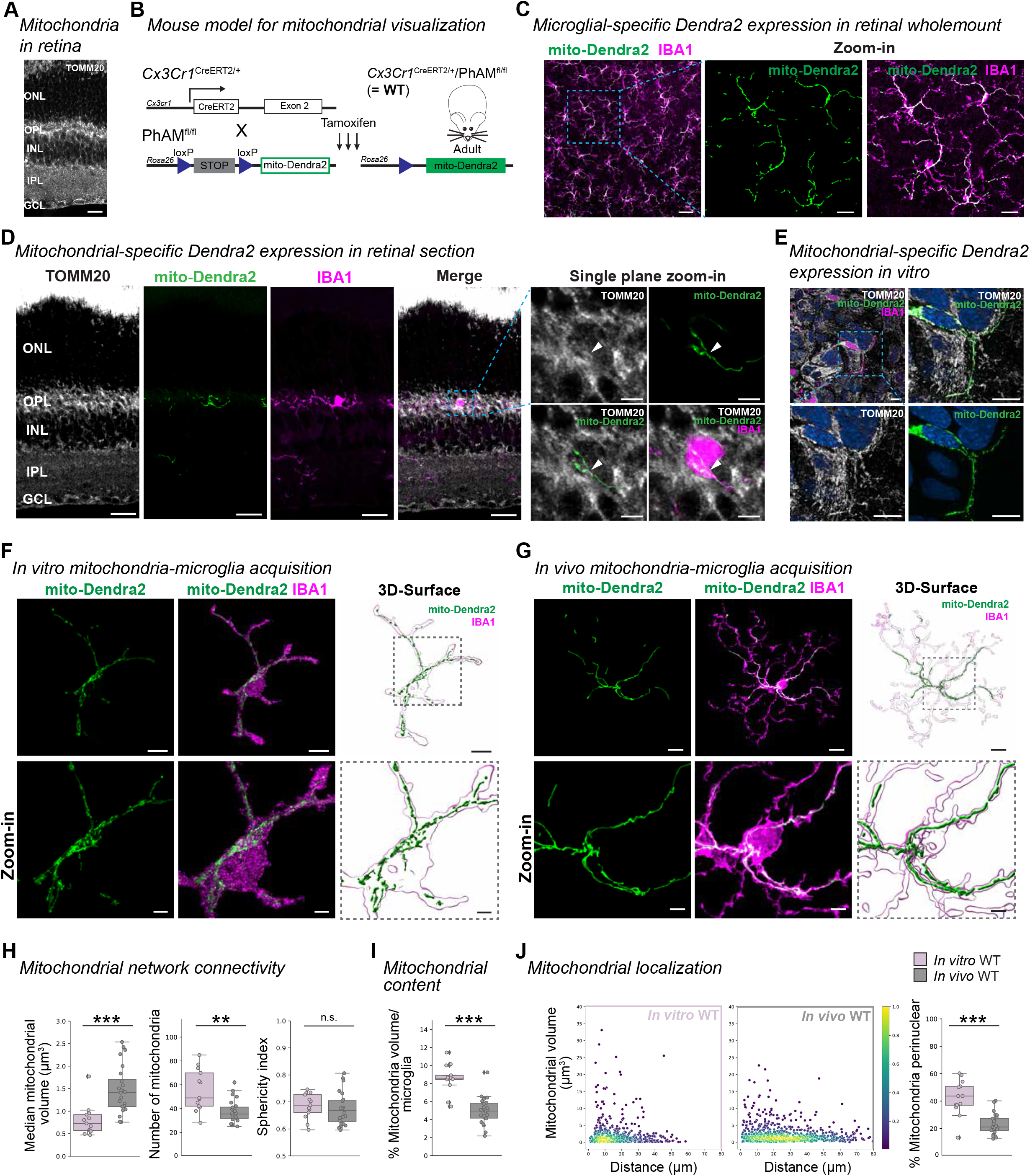
Mitochondria in microglia *in vivo* establish networks distinct from *in vitro*. (**A**) Retinal section immunostained for TOMM20 (white). Scale bar: 20 µm. ONL, outer nuclear layer. OPL, outer plexiform layer. INL, inner nuclear layer. IPL, inner plexiform layer. GCL, ganglion cell layer. (**B**) Schematic of mouse model for mitochondrial visualization with floxed mito-Dendra2 at the *Rosa26* locus (PhAM^fl/fl^). Pham^fl/fl^ are crossed to *Cx3cr1*^CreERT2^ for microglia-selective mito-Dendra2 expression (green) after receiving tamoxifen injection for three consecutive days. Referred to as WT throughout the study. (**C**) Overview image of immunostained microglia (IBA1, magenta) in the IPL expressing the mitochondrial label (Dendra2, green) in retinal wholemounts for WT. Scale bar: 50 µm, zoom-in: 20 µm. (**D**) TOMM20-immunostained retinal cross sections from WT mouse expressing mito-Dendra2 (green) and co-labeled with IBA1 (magenta). Scale bar: 20 µm. Single plane zoom-in (dashed outline) show overlap (white arrow) for mito-Dendra2 and TOMM20 (white). Scale bar: 5 µm. (**E**) TOMM20 (white) and IBA1 (magenta)-immunostained primary mixed glia culture with mito-Dendra2 (green) expression from WT mouse after 4-hydroxytamoxifen treatment. Scale bar: 15 µm. Dashed outline, zoom-in single plane. Blue: Hoechst labeled nuclei. Scale bar: 15 µm. (**F**-**G**) Immunostained microglia (IBA1, magenta) expressing mito-Dendra2 (green) in microglia *in vitro* from WT primary mixed glial cultures (**E**) or WT IPL *in vivo* (**F**). Scale bar: 10 µm. Next, 3D-surface rendering of IBA1 microglia (magenta outline) and Dendra2 mitochondria (green) with zoom-in (dashed line). Scale bar: 10 µm, zoom-in: 3 µm. (**H**-**J**) Mitochondrial parameters. Boxplot minimum and maximum: InterQuartile Range (IQR) around median (center line). Whiskers: 1.5 IQRs. Black diamond: outliers outside of 1.5 IQRs. Each overlaid point represents data point of a single microglia. *In vitro*: dusty pink. *In vivo*: grey. (**H**) Mitochondrial network connectivity determined by median mitochondrial volume (left, Wilcoxon rank sum test: *p* < 0.0001), number of organelles within a single cell (center, Welch’s t-test: *p* = 0.0033) and mean mitochondrial sphericity (right, t-test, *p =* 0.551017). (**I**) Percentage of mitochondrial volume per microglial volume. T-test: *p* < 0.0001. (**J**) Mitochondrial localization. Scatterplot depicting mitochondria volume *vs.* distance from the cell soma (0, origin) *in vitro* (left, dusty pink) or *in vivo* (right, grey) with pseudo-colored point density. Right: Percentage of total mitochondrial volume localized within the perinuclear region (0-10 µm). Wilcoxon rank sum test: *p* < 0.0001. ** p < 0.01, *** p < 0.001, ^n.s.^ p > 0.05: not significant. See **Supplementary Information** for experimental, retina and cell numbers, statistical tests and corresponding data.

Primary microglia in mixed glial culture differ in their morphological shape compared to microglia *in vivo* (**Figure 1F**-**G**). To evaluate the mitochondrial network in both conditions, we generated 3-dimensional (3D) surfaces of both mitochondria and microglia and performed volumetric analyses (**Figure 1F**-**G**, **S1A**, **Supplementary Video 1**-**2**). First, we quantified mitochondrial network connectivity for single cells in which we determined the median mitochondrial volume amongst all organelles of a cell (**Figure S1B**), the number of mitochondria, and the mean sphericity of mitochondria (**Figure S1C**). Microglia *in vitro* exhibited a reduced median mitochondrial volume and increased total number of mitochondria compared to *in vivo* (**Figure 1H**), indicating a more fragmented mitochondrial network, even though the sphericity did not differ. When we analyzed the mitochondrial content per cell (**Figure S1D**), we found that microglia *in vitro* had a significantly greater content compared to *in vivo* (**Figure 1I**), suggesting an imbalance of biogenesis and turnover between conditions, which can be associated with greater oxidative damage (Neurohr et al., 2021). To assess subcellular localization of mitochondria, we measured the distance from the centroid of each mitochondrion to the cell soma (**Figure S1E**) and represented the data in a scatter plot (**Figure S1F**). We found a greater percentage of a cell’s total mitochondrial volume was localized in the perinuclear region closer to the soma *in vitro* (**Figure 1J**, **S1G**). Collectively, these mitochondrial parameters demonstrate a difference between *in vitro* and *in vivo* conditions that align with a greater stress response in microglia *in vitro*. To substantiate this, we analyzed the expression of the endosomal-lysosomal marker CD68 (cluster of differentiation 68) in microglia (**Figures S2A**-**B**),which is upregulated in reactive microglia (Bauer et al., 1994; Damoiseaux et al., 1994). The CD68 volume was significantly greater *in vitro* and coincided with increased vesicle volumes distributed along the length of processes when compared to microglia *in vivo* (**Figure S2C**-**D**). Together, this data emphasizes that microglia *in vitro* are distinct from microglia *in vivo*, not only morphologically, but also at the mitochondrial level.

### Mitochondrial network alterations occur prominently in microglia proximal to dying neurons

To identify how mitochondrial networks of retinal microglia *in vivo* adapt to stress conditions such as neuronal death, we took advantage of the optic nerve crush (ONC) model (**Figure 2A**) (Levkovitch-Verbin et al., 2000; Li et al., 1999). Here, a mechanical injury at the retinal ganglion cell axons posterior to the optic disc induces their apoptosis (Li et al., 1999; Wohl et al., 2010), which we confirmed by an increased number of cleaved-caspase 3^+^/RBPMS^+^-retinal ganglion cells (**Figure 2B**). Microglia showed morphological adaptation to these environmental changes throughout the IPL after ONC (**Figure 2C**). At the single cell level, microglia increased expression of CD68 (**Figure S2A**-**C**), where vesicles were distributed throughout the length of the processes (**Figure S2D**). Accompanying this phenotype, microglia reduced their branching complexity, as verified by the increased number of Sholl intersections closer to the soma (**Figure S2E**). Next, we aligned mitochondrial network adaptations to this responsive microglial phenotype (**Figure 2D**-**E**). Relative to the naïve environment, microglia showed significantly reduced median mitochondrial volume and an increased number of mitochondria with greater sphericity after ONC (**Figure 2F**). Together with increased mitochondrial content (**Figure 2G**), and the subcellular localization of the mitochondrial population closer to the soma (**Figure 2H**), this data indicates a more fragmented mitochondrial network with altered content and localization in responsive IPL microglia.

**Figure 2.**
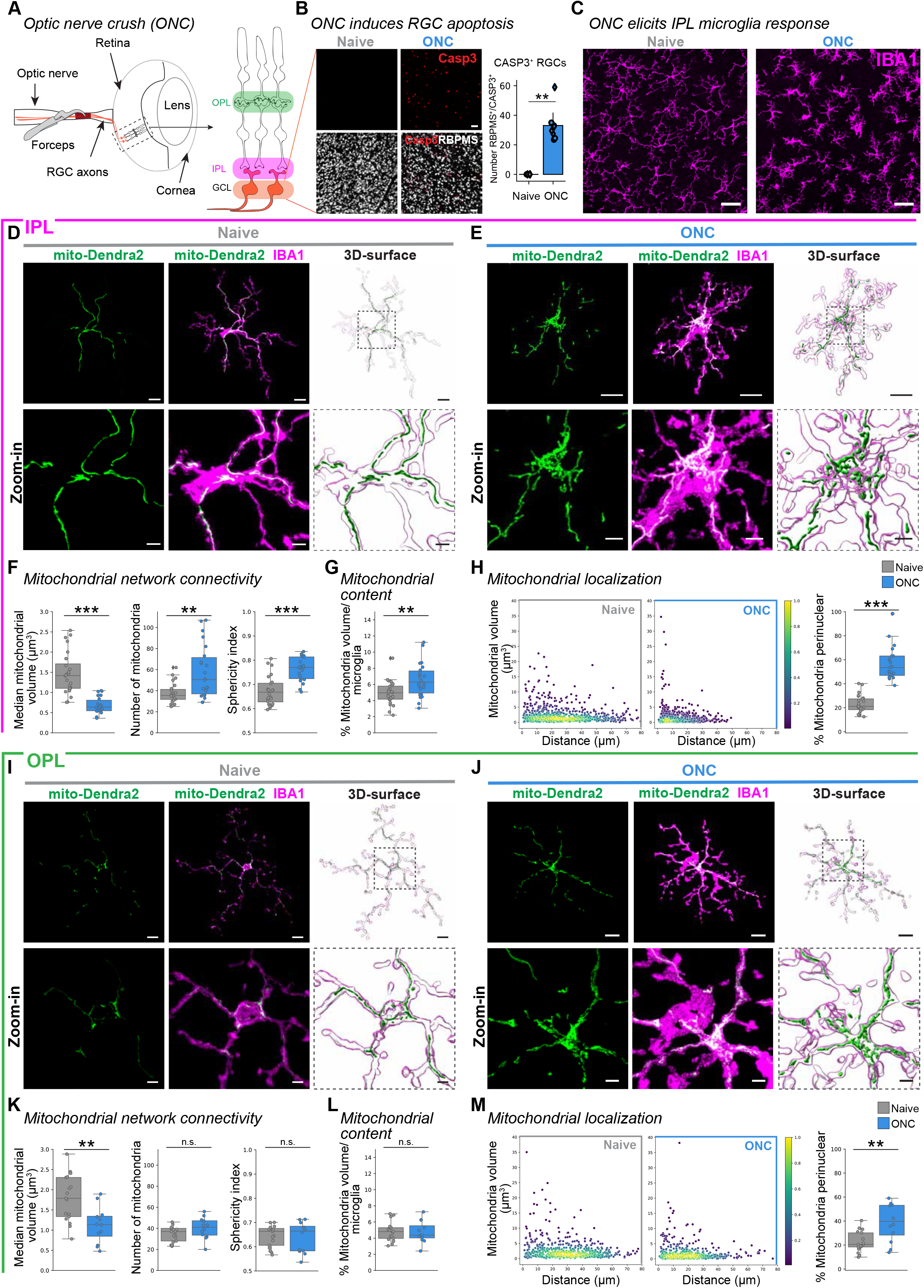
Regional differences in mitochondrial network connectivity, content and localization of ONC-responsive IPL and OPL microglia. (**A**) Schematic of optic nerve crush (ONC). Forceps induce unilateral damage (red) to axonal projections (orange) of retinal ganglion cells (RGCs) posterior to the optic disc. Right, retinal side-view schematic. OPL, outer plexiform layer. IPL, inner plexiform layer. GCL, ganglion cell layer. (**B**) Confocal images of RBPMS-immunostained naïve WT retinas or 5 days after ONC WT co-labeled with cleaved caspase-3 (CASP3, red) to identify dying retinal ganglion cells (RGCs, white). Right, bar plot indicate number of caspase-3^+^/RBPMS^+^ cells. Wilcoxon rank sum test: *p* = 0.0016. Scale bar: 50 µm. (**C**) Overview confocal image of immunostained IPL microglia (IBA1, magenta) in naïve WT (left) and 5 days after ONC WT (right) condition. Scale bar: 30 µm. (**D**-**M**) Comparison of mitochondrial response in naïve (magenta border) microglia WT (grey) and 5 days after ONC WT (blue). Boxplot minimum and maximum: InterQuartile Range (IQR) around median (center line). Whiskers: 1.5 IQRs. Black diamond: outliers outside of 1.5 IQRs. (**D**-**H**) Mitochondrial parameters for IPL microglia. (**D**-**E**) Mito-Dendra2 (green) expression in the IBA1-immunostained IPL microglia for naïve WT microglia (**D**) or ONC WT microglia (**E**). Next, corresponding 3D-surfaces. Below: Zoom-in of region of interest (dashed line) from image and 3D-surface. Scale bar: 10 µm, zoom-in: 3 µm. (**F**) Mitochondrial network connectivity determined by median mitochondrial volume (left, Welch’s t-test: *p* < 0.0001), number of organelles within a single cell (center, Wilcoxon rank sum test: *p* = 0.0018) and mean mitochondrial sphericity (right, T-test: *p* < 0.0001). (**G**) Percentage of mitochondrial volume per microglial volume. T-test: *p* = 0.0040. (**H**) Mitochondrial localization. Scatterplot depicting mitochondria volume *vs.* distance from the cell soma (0, origin) of the population of mitochondria in naïve WT (left) or ONC WT (right) microglia. Point density, pseudo-colored. Percentage of total mitochondrial volume localized within the perinuclear region. Wilcoxon rank sum test: *p* < 0.0001. (**I**-**M**) Mitochondrial parameters for OPL microglia. (**I**-**J**) Representative IBA1-immunostained OPL microglia (magenta) expressing mito-Dendra2 (green) from naïve WT (**I**) or ONC WT microglia (**J**). Next, corresponding 3D-surfaces. Below: Zoom-in of region of interest (dashed line) from image and 3D-surface. Scale bar: 10 µm, zoom-in: 3 µm. (**K**) Mitochondrial network connectivity determined by median mitochondrial volume (left, Student’s t-test: *p* = 0.0026), number of organelles within a single cell (center, Student’s t-test: *p* = 0.0900) and mean mitochondrial sphericity (Student’s t-test, *p* = 0.79389). (**L**) Percentage of mitochondrial volume per microglial volume. Student’s t-test: *p* = 0.8441. (**M**) Mitochondrial localization. Scatterplot depicting mitochondria volume *vs.* distance from the cell soma (0, origin) of the population of mitochondria in naïve WT (left) or ONC WT (right) microglia. Point density, pseudo-colored. Percentage of total mitochondrial volume localized within the perinuclear region. Welch’s t-test: *p* = 0.0045. ** p < 0.01, *** p < 0.001, ^n.s.^ p > 0.05: not significant. See **Supplementary Information** for retina and cell numbers, statistical tests and corresponding data.

To determine whether microglia residing more distant from the apoptotic retinal ganglion cells show a similar effect, we repeated the above analysis in microglia of the OPL. OPL microglia showed mild morphological adaptations (**Figure S3F**). Upon single cell analysis (**Figure 2I**-**J**), OPL microglia had significantly reduced median mitochondrial volume, whereas the number and the sphericity of mitochondria, and total content remained unaltered (**Figure 2K**-**L**). Similar to IPL microglia, the subcellular mitochondrial localization shifted towards the soma (**Figure 2M**), CD68 volume increased (**Figure S3G**-**J**), and the microglial branching complexity was moderately altered (**Figure S3K**). This data indicates that proximity to apoptotic neurons is a key determinant in the robustness of the mitochondrial-microglial response, where more distant microglia also exhibit some mitochondrial network alterations.

### Selective microglial UCP2^KO^ increases cellular and mitochondrial stress with no effect on the mitochondrial network

Electron leakage during oxidative phosphorylation leads to formation of mitochondrial superoxides and reactive oxygen species (ROS). Strategies to mitigate ROS include induction of mild uncoupling to increase the respiration rate and reduce electron leak, or detoxification of superoxides with cellular enzymes like superoxide dismutase 1 (SOD1) (West et al., 2011). However, these mitigation strategies are less effective in conditions of cellular damage or mitochondrial dysfunction, and can lead to ROS accumulation (Yu et al., 2020). One negative regulator of mitochondria ROS generation is uncoupling protein 2 (UCP2, Slc25a8), a mitochondrial-associated gene highly enriched in immune cells, namely retinal microglia (**Figure S4A**-**B**) (Bechmann et al., 2002; Brand and Esteves, 2005; Rupprecht et al., 2012; Siegert et al., 2012; Vozza et al., 2014; West et al., 2011). Upregulation of *Ucp2* transcript has been reported in microglia within disease environments (Hoang et al., 2020; O’Koren et al., 2019), and we confirmed increased *Ucp2* transcript expression in microglia five days after ONC (**Figure S4C**-**D**), making *Ucp2* a candidate gene to disrupt mitochondrial function and alter ROS accumulation.

Loss of UCP2 has been reported to elicit cellular ROS accumulation (Kizaki et al., 2002; Yasumoto et al., 2021). Thus, to evaluate this consequence on the mitochondrial network in retinal microglia, we generated microglia-specific UCP2^KO^ mice by crossing the *Ucp2*^fl/fl^ mouse with our mitochondrial-labeled WT mice (*Cx3cr1*^CreERT2/+^/PhAM^fl/fl^/*Ucp2*^fl/fl^, **Figure 3A**). This model, hereafter referred to as UCP2^KO^ mice, showed microglia-selective mitochondrial labeling (**Figure S4E**). We confirmed successful *Ucp2* knockout at both the transcript (**Figure S4F**) and protein level (**Figure S4G**-**I**) in which we performed qRT-PCR of FACS-isolated Dendra2^+^-retinal microglia three weeks after tamoxifen induction and Western blot analysis of microglia after 4-OHT (4-hydroxytamoxifen) treatment in primary mixed glial cultures, respectively. To determine the consequences of UCP2-knockout on microglial ROS levels, we first compared mitochondrial superoxide in retinal microglia from naïve WT and UCP2^KO^ mice using FACS-sorted MitoSOX stained cells. UCP2^KO^ mice exhibited a significantly increased percentage of MitoSOX^+^-microglia (**Figure 3B**). This coincided with significantly enhanced gene transcript expression of the detoxifying enzyme *Sod1* (**Figure 3C**), together indicating elevated mitochondrial and cellular ROS in UCP2^KO^ microglia. When we compared the mitochondrial networks of UCP2^KO^ to WT in the naïve condition, we did not detect differences in mitochondrial network connectivity, content, or localization (**Figure 3D**-**G**). Interestingly, the overall expression of CD68 was slightly elevated in UCP2^KO^ microglia (**Figure 3H**-**J**) and aligned with increased *CD68* transcript (**Figure 3K**), whereas the microglial morphology remained unaffected (**Figure 3L**). Together, this data indicates increased cellular stress in UCP2^KO^ microglia with no direct effect on the mitochondrial network in the naïve environment.

**Figure 3.**
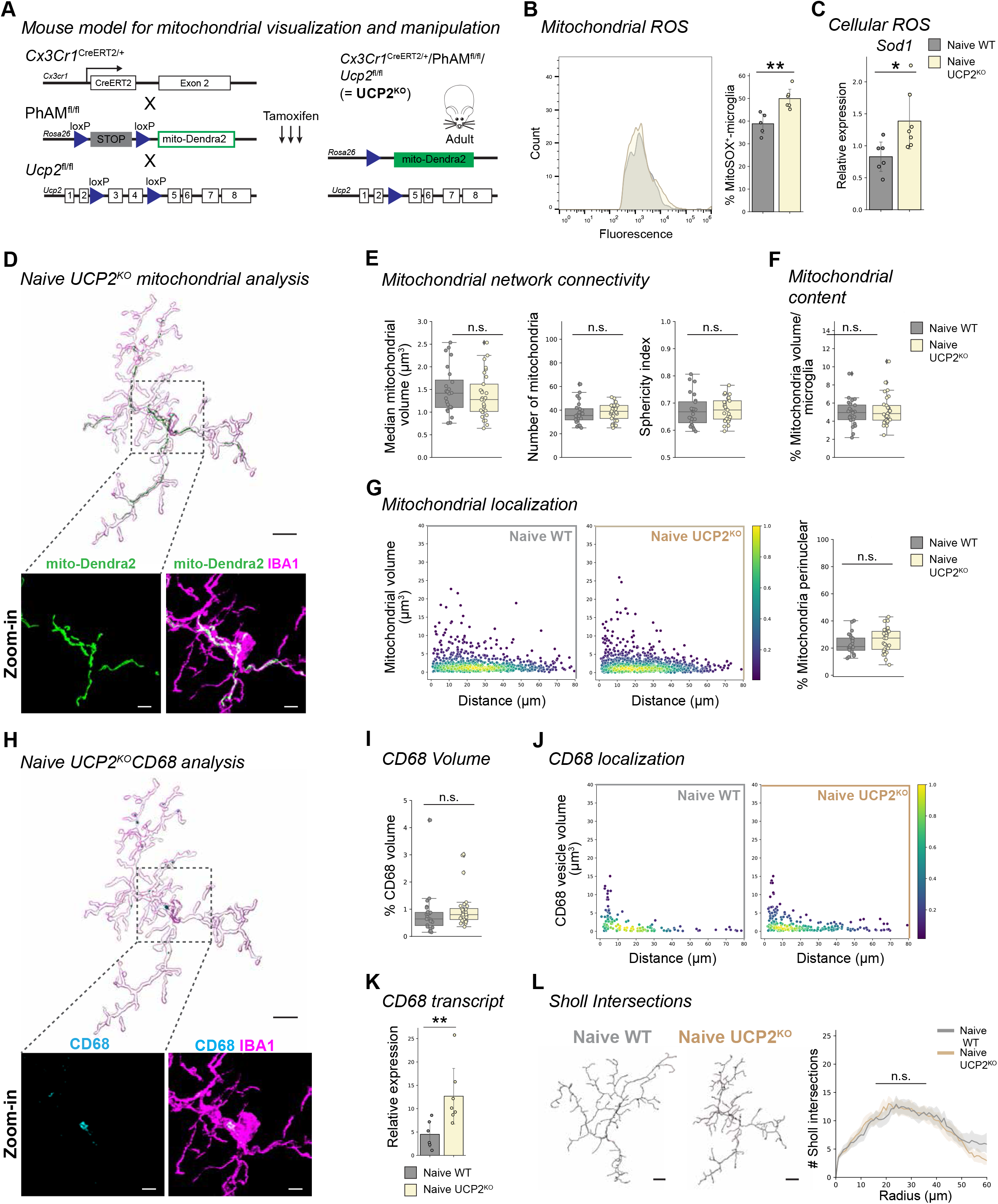
Mitochondrial networks are unaffected by increased stress from microglia-selective knockout of UCP2. (**A**) Schematic of mouse model. *Cx3cr1*^CreERT2/+^/Pham^fl/fl^/*Ucp2*^fl/fl^ for selective mitochondria-labeling in parallel with UCP2-knockout. Upon tamoxifen injection for three consecutive days in adult mice, *Cx3cr1*^CreERT2^-expressing microglia excise the stop cassette in Pham^fl/fl^ and exon 3 and 4 of *Ucp2* to induce selective mito-Dendra2 (green) expression and UCP2-knockout (here after referred as UCP2^KO^). (**B**) Frequency plot of MitoSOX fluorescence from FACSed mito-Dendra2^+^ retinal microglia for naïve WT (grey) and naïve UCP2^KO^ (tan). Corresponding bar plot of percentage of microglia that are MitoSOX^+^. Student’s t-test: *p* = 0.0033. (**C**) Bar plot depicting relative *Sod1* transcript expression from FACSed mito-Dendra2^+^ microglia in naïve WT or UCP2^KO^. Student’s t-test: *p* = 0.0141. (**D**) 3D-surface of representative IBA1-immunostained microglia (magenta) expressing mito-Dendra2 (green) from naïve UCP2^KO^. Below: Zoom-in of region of interest (dashed line) showing confocal images of mito-Dendra2 (green) expression or co-labeled IBA1-immunostaining (magenta). Scale bar: 10 µm, zoom-in: 3 µm. (**E**-**G**) Mitochondrial parameters. Boxplot minimum and maximum: InterQuartile Range (IQR) around median (center line). Whiskers: 1.5 IQRs. Black diamond: outliers outside of 1.5 IQRs. Each overlaid point represents data point of a single microglia. Naïve WT: grey. Naïve UCP2^KO^: tan. (**E**) Mitochondrial network connectivity determined by median mitochondrial volume (left, Student’s t-test: *p* = 0.4131), number of organelles within a single cell (center, Student’s t-test: *p* = 0.8337) and mean mitochondrial sphericity (right, Student’s t-test, *p* = 0.8732). (**F**) Percentage of mitochondrial volume per microglial volume. Wilcoxon rank sum test: *p* = 0.7410. (**G**) Mitochondrial localization. Scatterplot depicting mitochondria volume *vs.* distance from the cell soma (0, origin) of the population of mitochondria in naïve WT (left) or naïve UCP2^KO^ (right) microglia. Point density, pseudo-colored. Percentage of total mitochondrial volume localized within the perinuclear region. Student’s t-test: *p* = 0.1941. (**H**) 3D-surface of the IBA1-immunostained microglia (magenta) from Figure 3D co-labeled with CD68 (cyan). Below: Zoom-in of region of interest (dashed line) showing confocal images of CD68 (cyan) or co– labeled with IBA1-immunostaining (magenta). Scale bar: 10 µm, zoom-in: 3 µm. (**I**) Percentage of total CD68 volume per microglia volume. Boxplot minimum and maximum: InterQuartile Range (IQR) around median (center line). Whiskers: 1.5 IQRs. Black diamond points: outliers outside 1.5 IQRs. Wilcoxon rank sum test: *p* = 0.0847. (**J**) CD68 vesicle localization. Scatterplot depicting CD68 vesicle volume *vs.* distance from the cell soma (0, origin) of the population of vesicles in naïve WT (left) and naïve UCP2^KO^ (right) conditions. Point density, pseudo-colored. (**K**) Bar plot depicting relative *Cd68* transcript expression from FACSed mito-Dendra2^+^ microglia in naïve WT or naïve UCP2^KO^. Student’s t-test: 0.0062. (**L**) 3D-filament tracings of microglia from naïve WT (left) or naïve UCP2^KO^ (right). Scale bar: 10 µm. Line plot for mean number of Sholl intersections per radial distant from the soma (µm) with 95% confidence interval band. Linear mixed effects model: *p* = 0.9583. * p < 0.05, ** p < 0.01, ^n.s.^ p > 0.05: not significant. See **Supplementary Information** for retina and cell numbers, statistical tests and corresponding data.

### UCP2^KO^ allows microglial response and mitochondrial adaptations after ONC

Based on previous studies reporting that UCP2^KO^ prevents mitochondrial fission and microglial reactivity after high fat diet (Kim et al., 2019; Toda et al., 2016), we anticipated no mitochondrial fragmentation or microglial response in the ONC injury model. Unexpectedly, ONC UCP2^KO^ microglia showed increased CD68 volume, where vesicles were localized throughout the length of processes (**Figure 4A**-**D**), and reduced branching complexity (**Figure 4E**) compared to naïve UCP2^KO^. This aligned with the mitochondrial network becoming more fragmented as reflected in the reduced median mitochondrial volume and increased sphericity (**Figure 4I-K**). At the same time, the number of mitochondrial organelles surprisingly remained similar (**Figure 4H**, center) even though the mitochondrial content was increased (**Figure 4I**). This deviation in mitochondria number did not align with the expected changes for mitochondrial connectivity seen in WT ONC microglia (**Figure 2K**– **L**), indicating differences in UCP2^KO^ mitochondrial network adaptations.

**Figure 4.**
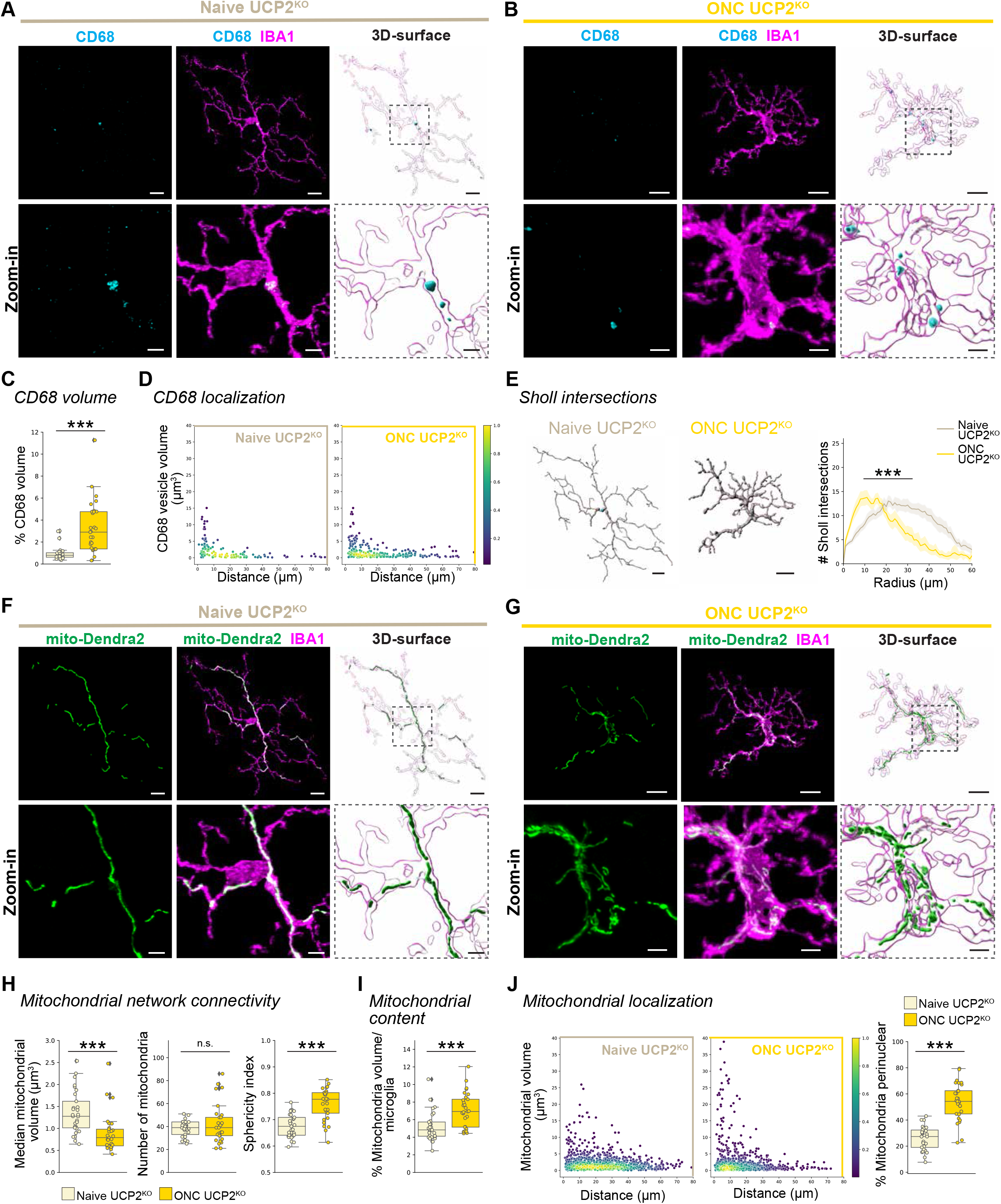
Mitochondrial alterations occur in responsive UCP2^KO^ IPL microglia after ONC. (**A-J**) Comparison of microglial and mitochondrial response in naïve UCP2^KO^ (tan) and 5 days after ONC UCP2^KO^ (gold). (**A**-**B**) Representative IBA1-immunostained microglia (magenta) co-labeled with CD68 (cyan) from naïve UCP2^KO^ (**A**) or ONC UCP2^KO^ microglia (**B**). Next, corresponding 3D-surfaces. Below: Zoom-in of region of interest (dashed line) from image and 3D-surface. Scale bar: 10 µm, zoom-in: 3 µm. (**C**) Percentage of total CD68 volume per microglia volume. Boxplot minimum and maximum: InterQuartile Range (IQR) around median (center line). Whiskers: 1.5 IQRs. Black diamond points: outliers outside 1.5 IQRs. Wilcoxon rank sum test: *p* < 0.0001. (**D**) CD68 vesicle localization. Scatterplot depicting CD68 vesicle volume *vs.* distance from the cell soma (0, origin) of the population of vesicles in naïve UCP2^KO^ (left) and ONC UCP2^KO^ (right, gold) conditions. Point density, pseudo-colored. (**E**) 3D-filament tracings of microglia from naïve UCP2^KO^ (left) or ONC UCP2^KO^ (right). Scale bar: 10 µm. Line plot for mean number of Sholl intersections per radial distant from the soma (µm) with 95% confidence interval band. Linear mixed effects model: *p* < 0.0001. (**F**-**J**) Mito-Dendra2 (green) expression in the IBA1-immunostained microglia from (**A**-**B**) for naïve UCP2^KO^ microglia (**F**) or ONC UCP2^KO^ microglia (**G**). Next, corresponding 3D-surfaces. Below: Zoom-in of region of interest (dashed line) from image and 3D-surface. Scale bar: 10 µm, zoom-in: 3 µm. (**H**-**J**) Mitochondrial parameters. Boxplot minimum and maximum: InterQuartile Range (IQR) around median (center line). Whiskers: 1.5 IQRs. Black diamond: outliers outside of 1.5 IQRs. (**H**) Mitochondrial network connectivity determined by median mitochondrial volume (left, Wilcoxon rank sum test: *p* < 0.0001), number of organelles within a single cell (center, Wilcoxon rank sum test: *p* = 0.930881) and mean mitochondrial sphericity (right, Student’s t-test: *p* < 0.0001). (**I**) Percentage of mitochondrial volume per microglial volume. Wilcoxon rank sum test: *p* = 0.0003. (**J**) Mitochondrial localization. Scatterplot depicting mitochondria volume *vs.* distance from the cell soma (0, origin) of the population of mitochondria in naïve UCP2^KO^ (left) or ONC UCP2^KO^ (right) microglia. Point density, pseudo-colored. Percentage of total mitochondrial volume localized within the perinuclear region. Student’s t-test: *p* < 0.0001. *** p < 0.001, ^n.s.^ p > 0.05: not significant. See **Supplementary Information** for retina and cell numbers, statistical tests and corresponding data.

### Loss of UCP2^KO^ elicits a sexually-dimorphic microglial response in ONC

Next, we looked more closely at the relationship between conditions (naïve, ONC) and genotypes (WT, UCP2^KO^) using principal component (PC) analysis. PC analysis allows unbiased visualization of six parameters for each analyzed microglia (**Figure S1, Methods**) and identification of patterns within the dataset. In naïve conditions, WT and UCP2^KO^ microglial profiles intermingled in the same PC space (**Figure 5A**), corroborating the results from **Figure 3**. Naïve and ONC microglial profiles separated along the first PC for both WT and UCP2^KO^, aligning the with microglial-mitochondrial response in **Figure 2** and **Figure 4**. However, the ONC microglial profiles showed less prominent intermingling between genotypes, where UCP2^KO^ showed greater spread along the second PC (**Figure 5A**). A recent study indicated that UCP2 loss in microglia results in anxiety phenotypes in male mice (Yasumoto et al., 2021), therefore we annotated sex as an additional factor in the PC space. Whereas sexes were intermingled for WT profiles in both naïve and ONC conditions (**Figure 5B**), UCP2^KO^ microglial profiles diverged along the second PC only after ONC (**Figure 5C**), suggesting a sexually-dimorphic response to ONC in UCP2^KO^ microglia.

**Figure 5.**
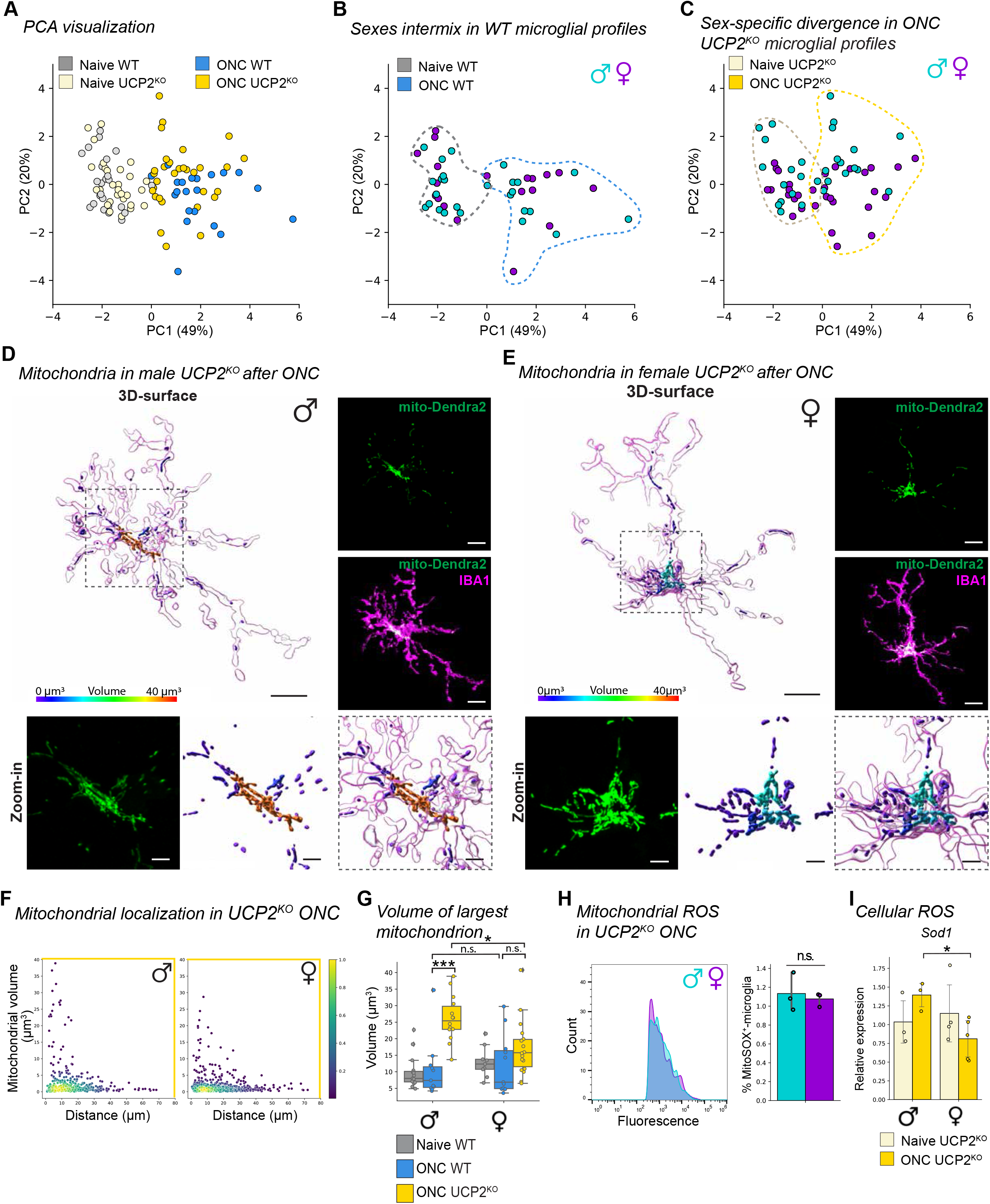
Male UCP2^KO^ microglia resort to mitochondrial hyperfusion for stress mitigation. (**A**-**C**) Principle component analysis (PCA) of microglia from WT and UCP2^KO^ in naïve and ONC conditions. The first two principle components (PC) are visualized. Each dot represents a single microglial profile defined by the following six parameters: microglial Sholl index, CD68 volume, median mitochondrial volume, number of mitochondria, % mitochondrial volume per microglia, % mitochondria perinuclear obtained from aforementioned genotype and condition (**Figure S1**). PC1 and PC2 describe 49% and 20% of the explained variance across the population, respectively. Each subpanel highlights the following comparisons: (**A**) Naïve microglial profiles for WT (grey) and UCP2^KO^ (tan) and ONC microglial profiles for WT (blue) and UCP2^KO^ (gold). (**B**-**C**) Naïve and ONC microglial profiles in male (turquoise) and female (purple) for WT (**B**) and for UCP2^KO^ (**C**). Dashed line: Reference of distributions shown in (**A**). (**D**-**E**) Representative 3D-surface renderings of IBA1 (magenta) and mito-Dendra2 (statistics-based volume spectral coloring) microglia from ONC-induced male UCP2^KO^ (**D**) and female UCP2^KO^ (**E**). Right: corresponding confocal images of IBA1-immonostained (magenta) microglia and mito-Dendra2 (green) expression. Below: Zoom-in of region of interest (dashed outline) of mito-Dendra2 (green) and corresponding 3D-surface with volume spectral coloring. Scale bar: 10 µm, zoom-in: 3 µm. Volume spectrum: 0 µm^3^ (blue) – 10 µm^3^ (red). (**F**) Mitochondrial localization. Scatterplot depicting mitochondria volume *vs.* distance from the cell soma (0, origin) of the population of mitochondria in ONC UCP2^KO^ male (left) or female (right) microglia. Point density, pseudo-colored. (**G**) Boxplot of largest mitochondrion per cell separated by sex. Boxplot minimum and maximum: InterQuartile Range (IQR) around median (center line). Whiskers: 1.5 IQRs. Black diamond: outliers outside of 1.5 IQRs. Naïve WT: grey. ONC WT: blue. ONC UCP2^KO^: gold. Kruskal-Wallis test: *p* < 0.0001. Selected Conover’s post-hoc comparisons with Holm p-adjustment: ONC ♂WT *vs.* ♂UCP2^KO^, *p* < 0.0001; ONC ♂WT *vs.* ♀WT, *p* = 1.0; ONC ♂UCP2^KO^ *vs.* ♀UCP2^KO^, *p* = 0.0387; ONC ♀WT *vs.* ♀UCP2^KO^, *p* = 0.1364. (**H**) Frequency plot of MitoSOX fluorescence from FACSed mito-Dendra2^+^ retinal microglia for ONC UCP2^KO^ male (turquoise) and female (purple). Corresponding bar plot of percentage of microglia that are mitoSOX^+^. Student’s t-test: 0.6707. (**I**) Bar plot depicting relative *Sod1* transcript expression from FACSed mito-Dendra2^+^ microglia in UCP2^KO^ naïve and ONC microglia. Student’s t-test: *p* = 0.0377. * p < 0.05, ^n.s.^ p > 0.05: not significant. See **Supplementary Information** for retina and cell numbers, statistical tests, other post-hoc comparisons and corresponding data.

### Male UCP2^KO^ microglia induce mitochondrial hyperfusion to mitigate stress

To identify which of the six parameters differed between sexes, we first referenced the PC loadings where the highest contributing factor along the second PC was mitochondrial content (**Figure S5A**). When separating the volumetric analyses by sex, UCP2^KO^ males showed significantly higher mitochondrial content per cell compared to female ONC UCP2^KO^ conditions (**Figure S5B**). Additionally, male ONC UCP2^KO^ exhibited a greater median mitochondrial volume and reduced mitochondrial number compared to female ONC UCP2^KO^ microglia, aligning with a less fragmented network (**Figure S5C**). To resolve these sex-based differences, we revisited the UCP2^KO^ microglial and mitochondrial images (**Figure 5D**-**E, Supplementary Video 3**) and separated the localization scatter plots. Here, we found that male ONC UCP2^KO^ microglia had a higher frequency of mitochondria with large volumes close to the cell soma relative to female ONC UCP2^KO^ (**Figure 5F**). When we compared the largest-volume mitochondrion per cell as a metric for highly-connected or hyperfused mitochondria, we found that male UCP2^KO^ had significantly greater volumes than female UCP2^KO^ or WT after ONC (**Figure 5G**), indicating a sexually-dimorphic hyperfused mitochondrial phenotype in UCP2^KO^ microglia.

Previously, mitochondrial hyperfusion has been described as a transient phenotype in stress-induced environments that protects cellular and mitochondrial health (Gomes et al., 2011; Tondera et al., 2009). To investigate whether the observed hyperfused mitochondrial networks occurred in response to differences in ROS levels between sexes, we quantified MitoSOX^+^-microglia after ONC in UCP2^KO^ for each sex. Males and females showed a comparable level of mitochondrial superoxide (**Figure 5H**), suggesting a similar production of intrinsic mitochondrial ROS. Interestingly, we detected opposing trends in *Sod1* transcript expression in UCP2^KO^ microglia after ONC, where males had significantly greater *Sod1* expression compared females after ONC (**Figure 5I**), indicating differences between sexes in the strategy for superoxide detoxification. This difference was more pronounced in WT microglia after ONC, where only males significantly increase *Sod1*, yet the production of mitochondrial superoxide remained similar in females (**Figures S5D**-**E**), suggesting sex-based differences in microglial ROS mitigation.

### Ovariectomized female UCP2^KO^ microglia evoke a male mitochondrial phenotype after ONC

Estrogens have been implicated to reduce oxidative stress and play a protective role in autoimmune and neurodegenerative conditions (Di Florio et al., 2020; Nakazawa et al., 2006; Razmara et al., 2007). Furthermore, both microglia and mitochondria express estrogen receptors (Baker et al., 2004; Yang et al., 2004). Since females did not implement stress-mitigation strategies *via* mitochondrial hyperfusion (**Figure 5E**-**G**) or increased *Sod1* transcription (**Figure 5I**, **S5D**), we evaluated whether circulating estrogens contribute to the sexually divergent mitochondrial phenotype in UCP2^KO^ after ONC. Thus, we repeated the experiment with ovariectomized UCP2^KO^ female mice (**Figure 6A**) and quantified the mitochondrial networks in naïve and ONC (**Figure 6B**). The mitochondrial connectivity in ovariectomized UCP2^KO^ female microglia exhibited decreased median mitochondrial volume, no change in mitochondria number, and significantly increased mitochondrial sphericity (**Figure 6C**). Furthermore, the mitochondrial content and their percentage in the perinuclear region increased after ONC (**Figure 6D**-**E**), together indicating mitochondrial adaptations to ONC in ovariectomized UCP2^KO^ females. Remarkably, the mitochondrial distribution in ovariectomized ONC UCP2^KO^ females reflected a similar increase in frequency of large-volume organelles close to the microglia soma (**Figure 6E**) as observed for UCP2^KO^ males (**Figure 5F**). When we analyzed the largest-volume mitochondrion per cell, UCP2^KO^ ovariectomized females had similarly large volumes as UCP2^KO^ males (**Figure 6F**).

**Figure 6.**
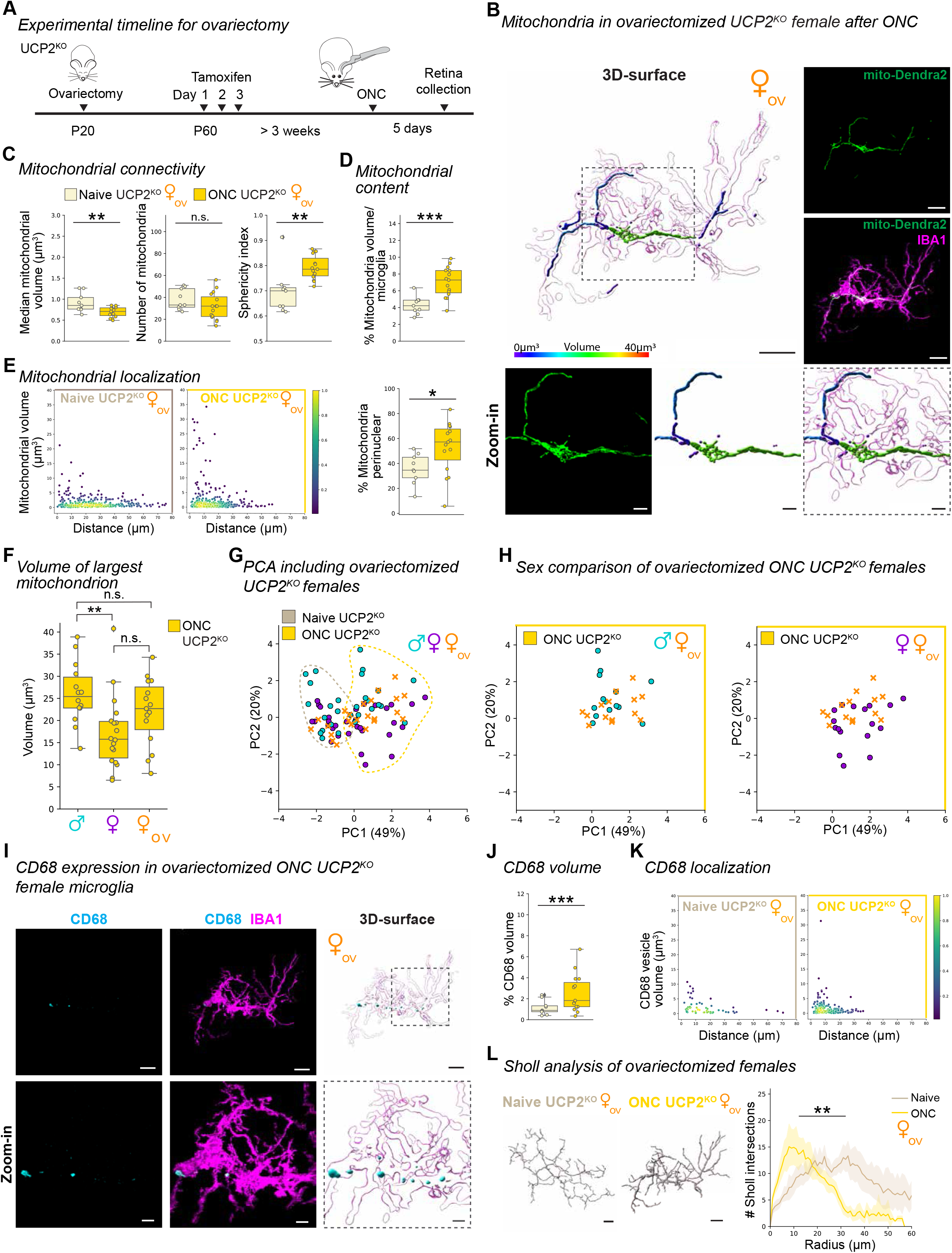
Ovariectomized female UCP2^KO^ microglia exhibit mitochondrial hyperfusion phenotype after ONC. (**A**) Experimental timeline. Ovariectomy performed at postnatal day 20 (P20) and three consecutive tamoxifen injections were administered daily for three days starting at P60. Optic nerve crush (ONC) was performed at least 3 weeks after tamoxifen injection and retinas collected five days after ONC. (**B**) Representative 3D-surface renderings of IBA1 (magenta) and mito-Dendra2 (statistics-based volume spectral coloring) from ONC-induced ovariectomized female UCP2^KO^ microglia. Right: corresponding confocal images of IBA1-immonostained (magenta) and mito-Dendra2 (green) expression. Below: Zoom-in of region of interest (dashed outline) of mito-Dendra2 (green) and corresponding 3D-surface with volume spectral coloring. Scale bar: 10 µm, zoom-in: 3 µm. Volume spectrum: 0 µm^3^ (blue) – 10 µm^3^ (red). (**C**-**G**) Comparison of mitochondrial response in ovariectomized females. Boxplot minimum and maximum: InterQuartile Range (IQR) around median (center line). Whiskers: 1.5 IQRs. Black diamond: outliers outside of 1.5 IQRs. Naïve UCP2^KO^: tan. ONC UCP2^KO^: gold. (**C**) Mitochondrial network connectivity determined by median mitochondrial volume (left, Student’s t-test: *p* = 0.0046), number of organelles within a single cell (center, Student’s t-test: *p* = 0.1939) and mean mitochondrial sphericity (right, Wilcoxon rank sum test: *p* = 0.0024). (**D**) Percentage of mitochondrial volume per microglial volume. Student’s t-test: *p* = 0.0004. (**E**) Mitochondrial localization. Scatterplot depicting mitochondria volume *vs.* distance from the cell soma (0, origin) of the population of mitochondria in microglia. Point density, pseudo-colored. Percentage of total mitochondrial volume localized within the perinuclear region. Student’s t-test: *p* = 0.0320. (**F**) Largest mitochondrion per cell separated by sex. One-way ANOVA: *p =* 0.0088. Selected Tukey’s post-hoc comparisons: ONC ♂UCP2^KO^ *vs.* ONC ♀UCP2^KO^, *p* = 0.0065; ONC ♂UCP2^KO^ *vs.* ONC ♀_ov_UCP2^KO^, *p* = 0.3297, ONC ♀UCP2^KO^ *vs.* ONC ♀_ov_UCP2^KO^, *p* = 0.1938. (**G**) Principle component analysis (PCA) of microglia from male, female and ovariectomized UCP2^KO^ microglia in naïve and ONC conditions. The first two principle components (PC) are visualized. Naïve and ONC UCP2^KO^ microglial profiles in male (turquoise), female (purple) and ovariectomized females (orange x-marker). Dashed outline: Reference of distributions shown in (Figure 5A). (**H**) Comparison of ovariectomized ONC UCP2^KO^ female (orange x-marker) microglial profiles with male (turquoise, left) or female (purple, right) ONC UCP2^KO^ microglial profiles. (**I**) Ovariectomized female ONC UCP2^KO^ IBA1-immunostained microglia (magenta) from Figure 6B co-labeled with CD68 (cyan) and corresponding 3D-surfaces. Below: Zoom-in of region of interest (dashed line) from image and 3D-surface. Scale bar: 10 µm, zoom-in: 3 µm. (**J**) Percentage of total CD68 volume per microglia volume in WT and UCP2^KO^ ovariectomized females in naïve and ONC conditions. Boxplot minimum and maximum: InterQuartile Range (IQR) around median (center line). Whiskers: 1.5 IQRs. Black diamond points: outliers outside 1.5 IQRs. Wilcoxon rank sum test: *p* < 0.0001. (**K**) CD68 vesicle localization. Scatterplot depicting CD68 vesicle volume *vs.* distance from the cell soma (0, origin) of the population of vesicles from ovariectomized females in UCP2^KO^ naïve (left) and UCP2^KO^ ONC (right) conditions. Point density, pseudo-colored. (**L**) 3D-filament tracings of microglia from UCP2^KO^ naïve (left) or ONC (right). Scale bar: 10 µm. Line plot for mean number of Sholl intersections per radial distant from the soma (µm) with 95% confidence interval band. Linear mixed effects model: *p* = 0.0018. *p < 0.05, ** p < 0.01, *** p < 0.001, ^n.s.^ p > 0.05: not significant. See **Supplementary Information** for retina and cell numbers, statistical tests and corresponding data.

To substantiate that ovariectomized ONC UCP2^KO^ females exhibit a mitochondrial phenotype similar to males, we represented all UCP2^KO^ microglial profiles in the PC space. As anticipated, the ovariectomized UCP2^KO^ females separated along the first PC into the microglial profiles for the respective naïve or ONC condition (**Figure 6G**). In the ONC condition, ovariectomized UCP2^KO^ females trended toward males along the second PC (**Figure 6H**). Aligning with the ONC-responsive mitochondrial phenotype, ovariectomized UCP2^KO^ females showed increased CD68 volume with greater distribution and reduced branching complexity (**Figure 6I-L**). Together, this data illustrates that circulating estrogens play a role in maintaining mitochondrial networks in female microglia, and are a contributing factor to the sexually-divergent stress-mitigation strategies in UCP2^KO^ microglia.

## Discussion

Our results provide a new perspective of mitochondrial networks in microglia *in vivo*, and demonstrate their adaptation to an injury environment and increased cellular stress. Importantly, we established sex-based differences in microglial stress mitigation strategies, which we uncovered at the mitochondrial network level upon UCP2 knockout. Extensive *in vitro* studies have shown that cellular energy demands of immune cells critically depend on balanced mitochondrial networks equilibrated by mitochondrial fission and fusion (Rambold and Pearce, 2018). Insight into these dynamics in microglia *in vivo* are mostly unknown due to technical limitations. Tissue-wide mitochondrial immunostaining requires advanced image segmentation techniques to decipher which mitochondria are within or outside the microglia (Erny et al., 2021). This necessitates high resolution images and experience in how to determine cell edges to avoid user-biases (Costa and Cesar, 2009). In the retina, we performed high resolution imaging of TOMM20 immunostaining and were challenged to perform image segmentation of microglial mitochondria for both the retina (**Figure 1D**), and in 2D-mixed primary glial culture (**Figure 1E**). In contrast, our mouse model allowed clear image segmentation of microglial mitochondria (**Figure 1C, F**-**G**). Alternative techniques like electron microscopy can be used to assess cross-sectional mitochondrial areas (Kim et al., 2019; Toda et al., 2016), however resolving the entire mitochondrial network of an intact microglia would require time-and labor-intensive serial sectioning followed by image segmentation (Bolasco et al., 2018). Our mouse model visualizes the mitochondrial network of an entire microglia *in vivo* and allowed us to resolve elongated, tubular organelles in naïve condition, which was absent *in vitro* (**Figure 1F**-**G**) (Nair et al., 2019; Park et al., 2013). Our results in ONC-responsive microglia *in vivo* corroborated a study reporting increased mitochondrial mass in responsive microglia isolated from brain tissue (Erny et al., 2021), indicating that results *in vivo* are achievable without microglial isolation and importantly, maintain the spatial and morphological information of the cell. To this end, we identified regional differences in mitochondrial network adaptations between microglia localized in the IPL and OPL after ONC, which are not attainable in cell-dissociation or *in vitro* studies (**Figure 2**, **S3**). Together, this emphasizes that our mouse model provides a valuable strategy for assessing the mitochondrial network of microglia in a region-specific manner without cell-isolation.

Mitochondrial networks are most commonly assessed using connectivity parameters, including organelle volume, number, and sphericity (Chaudhry et al., 2020). We found that these three parameters reliably identified fragmented networks in ONC-responsive IPL microglia (**Figure 2**), which was also the case when two of the three parameters were altered, highlighting the importance of using more than one metric to evaluate mitochondrial connectivity in the 3D environment. As an additional parameter, we also assessed the subcellular localization of each mitochondrion within individual microglia. Previous studies have shown that responsive microglia enhance their transcriptional activity (Hammond et al., 2019), a cellular feature facilitated by perinuclear localization of mitochondria (Al-Mehdi et al., 2012). We consistently detected an increased fraction of the mitochondria in the perinuclear region across genotypes and sexes after ONC (**Figure 2**, **4**, **6**, **S3**). Interestingly, this localization was one of only two mitochondrial parameters that significantly changed in OPL microglia (**Figure S3F**). The second parameter was decreased median mitochondrial volume (**Figure S3D**), which is indicative of mitochondrial fragmentation. Thus, both parameters suggest initial mitochondrial adaptations in microglia after ONC in the OPL niche more distant from the injury site. The combination of subcellular localization and the mitochondrial content per cell were critical to identify the hyperfused mitochondrial network (**Figure 5**). Hyperfused mitochondrial networks have been described *in vitro* as either a stress-induced pro-survival mechanism in conditions of nutrient deprivation or as a result of non-functional fission machinery (Abdullah et al., 2022; Frank et al., 2001; Gomes et al., 2011; Mitra et al., 2009; Qian et al., 2012; Tondera et al., 2009). We found that this phenotype also occurs in microglia *in vivo,* which we identified with significant increases in large-volume mitochondria (**Figure 5F-H**). This metric was most informative for identifying hyperfusion in microglia since the unique, 3D morphology of microglia makes it difficult to use a subjective classification method without quantification to identify hyperfused networks, which has been the common methodology for defining mitochondrial hyperfusion in *in vitro* studies (Gomes et al., 2011; Mitra et al., 2009; Tondera et al., 2009).

Our approach to evaluate the effects of increased cellular stress on mitochondrial networks *in vivo* was to endogenously manipulate stress levels through selective-knockout of UCP2, a negative regulator of ROS. UCP2^KO^ microglia showed increased intrinsic stress and CD68 expression in the naïve environment (**Figure 3B**-**C**, **I**-**K**), aligning with previous literature (Kim et al., 2019; Yasumoto et al., 2021). Although these stress levels were increased, we did not detect any significant mitochondrial network alterations in the naïve environment (**Figure 3**). It is possible that mitochondrial adaptations occurred earlier after knockout. However, our model limited us to a three week timepoint after UCP2-depletion when circulating Cx3Cr1^+^-monocytes have repopulated (Goldmann et al., 2013). This ensures that UCP2-knockout and mitochondrial-labeling is restricted to the resident microglia and thus confirms that our mitochondrial analysis is exclusive to this population in the event of monocyte infiltration after ONC injury. Nevertheless, we did detect mitochondrial fragmentation and a microglial response in the UCP2^KO^ ONC environment (**Figure 4I**, **J**-**L**), which was surprising since previous studies reported that UCP2 loss prevented these changes in the hypothalamus by assessing cross-sectional area and number of mitochondria in electron microscopy preparations (Kim et al., 2019; Toda et al., 2016). On the other hand, *Kim et al*.(Kim et al., 2019) reported prevention of mitochondrial fragmentation only in male mice, while females were largely unaffected in diet-induced obesity paradigms and their mitochondrial phenotypes were not reported. Thus, our results align with these previous reports and refine that UCP2 loss affects mitochondrial networks in a sex-specific manner, such that UCP2^KO^ induces hyperfusion in male microglia after ONC, while female UCP2^KO^ microglia still exhibit mitochondrial fragmentation.

Since male and female microglia responded differently to increased stress via UCP2^KO^ (**Figure 5**, **S5**), we suspect that estrogens may explain the difference in the female response. Circulating estrogens provide a protective role for neurons under oxidative stress or in injury conditions (Nakazawa et al., 2006; Zhang et al., 2009) and enhance antioxidant gene expression in female mitochondria (Borrás et al., 2003; Pinto and Bartley, 1969). Indeed, in absence of circulating estrogens, both male and ovariectomized female microglia rely on the same mechanism of mitochondrial hyperfusion to mitigate excess stress from UCP2^KO^ (Gomes et al., 2011; Tondera et al., 2009). On the other hand, estrogen may not be the only mechanism. In the absence of circulating estrogens *in vitro*, female but not male neurons were shown to utilize lipids as a pro-survival fuel source (Du et al., 2009), and in a separate study, it was shown that female microglia support protection from diet-induced obesity by an estrogen-independent increase in CX3CR1 signaling (Dorfman et al., 2017). This suggests that sex-differences in microglial stress responses may be driven by a combination of circulating estrogens and dimorphisms in transcriptional programs (Baker et al., 2004; Thion et al., 2018; Villa et al., 2018; Yang et al., 2004).

In conclusion, our study highlights the substantial importance of sex-mediated effects in microglia which are reflected in the mitochondrial network, and provides a foundation for mitochondrial network analysis of microglia *in vivo*.

## Author contributions

M.M. and S.S. conceived and developed experimental design collaboratively. M.M. wrote the initial draft with subsequent editorial input from S.S. M.M. performed experiments, created and implemented python data import for 3-dimensional analysis, performed statistical analysis, and created figures. M.M., F.S. and E.E.P. conceived and verified UCP2 protein knockout. G.C. performed FACS-based RT-qPCR experiments, and data analysis. F.S.U. and M.M. performed FACS-based MitoSOX experiments and data analysis. A.V. performed ovariectomy surgeries. All authors provided comment and discussion on the manuscript and approved the final version.

## Acknowledgements

We thank the Scientific Service Units (SSU) of ISTA through resources provided by the Imaging and Optics Facility (IOF), the Lab Support Facility (LSF), and the Pre-Clinical Facility (PCF) team, specifically Sonja Haslinger and Michael Schunn for excellent mouse colony management and support. This research was supported by the FWF Sonderforschungsbereich F83 (to E.E.P). We thank Bálint Nagy, Ryan John A. Cubero, Marco Benevento and all members of the Siegert group for constant feedback on the project and manuscript.

## Materials and Methods

### Animals

Animal housing and procedures were approved by the “Bundesministerium für Wissenschaft, Forschung und Wirtschaft (bmwfw) Tierversuchsgesetz 2012, BGBI. I Nr. 114/2012 (TVG 2012) under the number GZ: 2021-0.607.460. Mice were housed in a 12 hour light-dark cycle in individually ventilated cages in a controlled environment (room temperature 22 ± 1 °C; relative humidity 55 ± 10 %) with access to standard diet (rat/mouse maintenance diet (V1534-300) or mouse breeding diet (V1124-300), ssniff Spezialitäten GmbH) and autoclaved water provided *ad libitum* in the ISTA Preclinical facility. Founder mouse strains were obtained from Jackson Laboratories and backcrossed to C57Bl6/J (#000664) background for at least 10 generations: *Cx3cr1*^CreERT2^ (#020940), PhAM^fl/fl^ (#018385, (Pham et al., 2012)), *Ucp2*^fl/fl^ (#022394, (Kong et al., 2010)). Throughout the manuscript WT refers to *Cx3cr1*^CreERT2/+^/PhAM^fl/fl^, while UCP2^KO^ refers to *Cx3cr1*^CreERT2/+^/PhAM^fl/fl^/*Ucp2*^fl/fl^. In addition to our confirmation of the mito-Dendra2 localization at the mitochondria (**Figure 1D**-**E**), the original study verifies this localization in the mouse model (Pham et al., 2012). For UCP2 protein knockout primary microglia culture experiments (**Figure S4G**-**I**) mice did not contain the Dendra2 mitochondria label, and instead were *Cx3cr1*^CreERT2/+^ and Cx3cr1^CreERT2/+^/UCP2^fl/fl^.

### Tamoxifen administration for Cre-induced recombination

Tamoxifen (Sigma Aldrich, T5648-5G) was dissolved in corn oil (Sigma Aldrich, C8267-500ML) at a concentration of 20 mg/mL and sonicated for 40 minutes. Freshly prepared tamoxifen was administered via intraperitoneal injection daily (150mg/kg) for three days. The animals did not show any abnormal behavior or discomfort after gene knockout. All experimental groups (WT and UCP2^KO^ naïve and ONC) were tamoxifen-injected for two reasons; first, expression of the Dendra2 mitochondria label is only possible after Cre recombinase excision of the stop cassette and second, *Cx3Cr1*^CreERT2^ reporter lines have been shown to include some ‘leakiness’, *i.e.* tamoxifen-independent *Cre* recombination (Van Hove et al., 2020). *Cx3Cr1*^CreERT2^ mice also express CreERT2 in monocyte populations, therefore to ensure no interference from UCP2^KO^ circulating monocytes, ONC experiments were performed at least three weeks after tamoxifen induction when monocyte populations have turned over (Goldmann et al., 2013). This also provided sufficient time for adaptation of microglia to the UCP2 loss conditions and clearance of all tamoxifen metabolites (Jahn et al., 2018).

### Optic nerve crush (ONC) procedure

Mice were anesthetized in an induction chamber with 5% (v/v) isoflurane (Zoetis) supplied with oxygen at a flow rate of 0.6 L/min. After lack of a foot pinch reflex, mice were maintained at 2.5% (v/v) isoflurane applied through a nose cone while on a heating pad to maintain body temperature at 37°C. Proparacaine hydrochloride 0.5% ophthalmic eye drops (Ursapharm Arzneimittel GmbH) were applied to numb the eyes, and subcutaneous injection of 5mg/kg Metacam alleviated pain (Meloxacam, Boehringer Ingelheim). The lateral canthus was de-vascularized by clamping with a hemostat (Fine Science Tools) for 10 seconds. Using a Leica dissection microscope, a lateral canthotomy allowed visualization of the posterior pole. While firmly holding the conjunctiva with a jeweler forceps, the conjunctiva was cut perpendicular to the posterior pole. The surrounding muscle was carefully dissected as to not puncture the vascular plexus. The optic nerve was pinched 1mm from the posterior pole for 4 seconds using a curved N7 self-closing forceps (Dumont). Triple antibiotic ointment was applied to the eye directly after the surgery to prevent infection.

### Ovariectomy

*Cx3cr1*^CreERT2/+^/PhAM^fl/fl^ or *Cx3cr1*^CreERT2/+^/PhAM^fl/fl^/*Ucp2*^fl/fl^ prepubescent postnatal day 20 female mice were anesthetized using 5% (v/v) isofluorane supplemented with oxygen at a flow rate of 0.6L/min in an induction chamber (Hoffmann, 2018). Mice were maintained in 2% (v/v) isoflurane with the same flow rate after absence of a foot pinch reflex. Above the lumbar spine, the skin was exposed by shaving the fur using an electric razor, then sterilized with 70% (v/v) ethanol. A 1cm midline incision on the lower back allowed gentle dissection of the subcutaneous tissue to expose the muscular fascia and ovarian fat pad. A small incision into the peritoneal cavity for entry exposed the fallopian tube, which was used as a guide to identify the ovary. The ovary was removed via cauterization of the oviduct and blood vessels to prevent bleeding. The muscular fascia was sutured after the fallopian tube was replaced into the peritoneal cavity. After repeating the procedure to remove the contralateral ovary, the midline incision of the skin was sutured. Mice received subcutaneous injection of Metamizol (Sanofi Avenis, 200 mg/kg) during the surgical procedure and meloxicam (Boehringer-Ingelheim, 5mg/kg) to alleviate pain.

### Primary glial culture

Mixed glia culture were prepared from the published protocol by *Bronstein et al*. (Bronstein et al., 2013). Cortices from 3-5 murine pups aged P0-P2 were dissected under a sterile ventilation hood in ice-cold Hank’s buffered saline (ThermoFisher, #14025-050) and digested in 0.05% (v/v) Trypsin + 1X EDTA (ThermoFisher, #25300-054) for 15 minutes at 37°C. The trypsinization was stopped with the addition of serum-containing medium (DMEM, 10% (v/v) heat-inactivated FBS, 1% (v/v) Penicillin/Streptomycin, 1% (v/v) Non-essential amino acids (Sigma-Aldrich, M7145-100ML)). Cells were pelleted at 500×g for 5 min, washed one time with serum-containing medium, then passed through a 40µm cell strainer (Szabo-Scandic, #352340). For Western blotting, the resulting cell suspension was plated onto T75 cell culture flasks (TPP, #Z707503). The culture medium was replaced at day 3 and day 10.

### Primary microglia isolation

Two T75 flasks containing mixed primary glia cultures prepared in tandem were treated with 1µM 4-hydroxy tamoxifen (Sigma-Aldrich, SML1666-1ML) or vehicle at day 10. Three days later, microglia were isolated using mild trypsinization (0.8mM CaCl_2_ in 1×Trypsin-EDTA, ThermoFisher #25300-054) is added to the flask after a washing step with pre-warmed 1X PBS to remove inhibiting serum, then incubate at 37°C for 10-20 minutes, or until the non-microglial cells have just lifted from the flask forming a floating layer. The floating cells were collected with the supernatant, spun down at 10,000×g, washed once with 1X PBS, pelleted and snap-frozen. The microglia remaining on the flask were overlaid with 5ml 1X PBS, removed using a cell scraper, pelleted, and snap-frozen.

### Western blotting

Total cellular protein isolation, production of recombinant UCP2 in *E. coli*, and western blot (WB) was performed as described previously (Rupprecht et al., 2012, 2014). For WB analyses 10 µg of total protein were loaded on the SDS-gel, 1 ng of recombinant UCP2 was used as positive control for anti-UCP2 antibody detection. To ensure blotting homogeneity to nitrocellulose membranes, Ponceau S staining was performed. To relate UCP2 expression to loading control, the membranes were stripped, re-blocked and incubated with respective antibodies. The affinity purified antibody against the N-terminal sequence of mouse UCP2 (VGFKATDVPPTATVKF) was designed and evaluated in the Pohl laboratory (Rupprecht et al., 2012). Peptide synthesis, rabbit immunization and affinity purification were performed by PINEDA Antibody-service GmbH (Berlin, Germany). All other antibody source and concentrations are provided in **Table S2**. The production of recombinant murine UCP2, used as positive control, was performed as described in (Žuna et al., 2021).

### Retina sample preparation for fluorescence-activated cell sorting (FACS)

Adult animals expressing the mito-Dendra2 tag were anesthetized with isoflurane and decapitated. Retinas were immediately explanted and dissected in 1X PBS on ice. Each retina was transferred to a 1.5ml Eppendorf low-adhesion tube filled with the 800µl digestion buffer (1:8:1 Cysteine/EDTA solution (2.5mM Cysteine, 0.5mM EDTA (ethylenediaminetetraacetic acid) in HBSS, 10mM HEPES (4-(2-hydroxyethyl)-1-piperazineethanesulfonic acid) in HBSS, and 10mg/ml Papain, Roche #10108014001), incubated at 37°C for 10 minutes, then centrifuged for 2.5 min at 240×g. Samples were transferred on ice, the supernatant was discarded and 800µl 1mM EDTA in HBSS + 2% (v/v) FBS added. After two washes with 1mM EDTA in HBSS + 2% (v/v) FBS, the digested tissue was triturated 10-15 times with a pulled glass pipette, then filtered through a 70 µm strainer. For the measurement of mitochondrial reactive oxygen species, 5 µM MitoSox (ThermoFisher, #M36008) in HBSS was added to the digested tissue and pipetted to break apart the tissue prior to a 10 minute incubation at 37°C, then the sample was centrifuged for 3 minutes at 304×g at 4°C. Samples were carefully washed two times with 1mM EDTA in HBSS + 2% (v/v) FBS, triturated 10-15 times with a pulled glass pipette, then filtered through a 70 µm strainer.

### Fluorescent activated cell sorting (FACS)

FACS was performed using a SONY SH800SFP equipped with a 100 µm nozzle and sorting speed of approximately 8,000 events per second in normal purity mode. Microglia were gated from the population by setting the forward and side scatter and by the fluorescence of the mito-Dendra2 tag. From each retina, 100 microglia were sorted and immediately collected in one well of a 96-well plate (Eppendorf) filled with 5 µl cold lysis buffer from the NEBNext® Single Cell/Low Input cDNA Synthesis & Amplification Module (New England Bio Labs, #E6421L). After the sorting, the plate was shortly spun down to ensure all cells were collected at the bottom of the well and was immediately processed for cDNA synthesis. For MitoSOX based flow cytometry approximately 500,000 events were acquired for each experimental condition. All collected data were gated based first on forward and side scatter, and then on the fluorescence of the mito-Dendra2 tag and the MitoSOX. Data were analyzed using FlowJo ™ v10.8 Software (BD Life Sciences).

### cDNA synthesis

Sorted microglia were processed with the NEBNext® Single Cell/Low Input cDNA Synthesis & Amplification Module (New England Bio Labs, #E6421L) according to the manufacturer’s protocol.

### Reverse transcription quantitative real-time PCR and gene expression analysis

Primers (**Table S1**) were designed with the free PrimerQuest Tool from Integrated DNA Technologies (https://eu.idtdna.com/PrimerQuest/Home/Index). To ensure the target sequence of each primer, primers were blasted (https://www.ncbi.nlm.nih.gov/tools/primer-blast/). Their self-complementarity and folding probability were investigated using the UNAFold Web server (http://www.unafold.org/). Primer efficiencies were validated from the slope of four to five serial 1:4 dilutions of cDNA template according to equation (1). Primers with efficiencies between 90-110% were used.

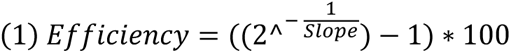

For gene expression analysis, RT-qPCR (Luna® Universal qPCR Master Mix; New England BioLabs; M3003L) was performed in 384-well plates (Bio-Rad; HSR4805) on a Roche Lightcycler 480 according to the manufacturer’s manual. Total reaction volume was 10μl containing 1μl of 1:10 diluted cDNA as template and a final concentration of 0.25μM for each primer. Cycle conditions were 60 seconds at 95°C for initial denaturation, followed by 40 cycles of denaturation (15 seconds; 95°C) and annealing/extension (30 seconds; 60°C).

Each run was completed with a melting curve analysis to confirm amplification of only one amplicon. Each PCR reaction was run in triplicate from which a mean Cq value was calculated and used for further analysis.

### Analysis of RT-qPCR results

Fold change differences between each condition and WT_naive_ were calculated according to delta-delta Ct method (Schmittgen and Livak, 2008). dCq values were obtained by normalizing mean Cq values to the reference housekeeping gene (GAPDH) measured within the same experiment (Equation 2). ddCq values were then calculated by normalizing dCq values to the respective control condition (WT_naive_) within each experimental repetition (Equation 3). Fold changes were obtained by transforming ddCq values from log2-scale to linear scale (Equation 4). These fold changes were used for data visualization. Exclusion criteria were based on IBA1 and GFAP fold change expression to determine purity of cell isolation, which would indicate substantial contamination from astrocytes and Müller glia.

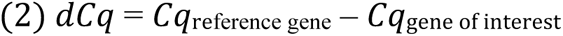

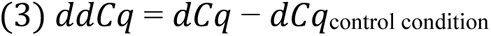

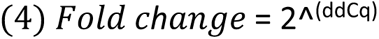

### Retina dissection and fixation

Animals were briefly anesthetized with isoflurane (Zoetis) until breathing slowed, then cervical dislocation was performed. Retinas were immediately dissected from the enucleated eyes in 1X phosphate buffered saline (PBS) and fixed in 4% (w/v) paraformaldehyde for 30 minutes. After 3× PBS washes, retinas were put in 30% (w/v) sucrose in 1X PBS overnight at 4°C for either –80°C freezer storage or immunostaining.

### Immunostaining

Staining was carried out in 24-well tissue culture plates (TPP, #92024) always protected from light to preserve the endogenous mito-Dendra2 fluorophore. Retinas saturated in sucrose (30% sucrose (w/v) in 1X PBS) underwent three freeze/thaw cycles using dry ice, then were washed 3× with 1X PBS. Next, a blocking step using blocking solution (1% (w/v) bovine serum albumin (Sigma A9418), 5% (v/v) Triton X-100 (Sigma T8787), 0.01% (v/v) sodium azide (VWR 786-299), and 10% (v/v) goat (Millipore S26) or donkey (Millipore S30) serum) was carried out on an orbital shaker for 1 hour at room temperature. Primary antibodies (**Table S2**) were diluted in antibody solution (1% (w/v) bovine serum albumin, 5% (v/v) triton X-100, 0.01% (v/v) sodium azide, 3% (v/v) goat or donkey serum) and incubated for three nights at 4°C on an orbital shaker. Following 3× PBS washes for 30 mins each, secondary solutions were prepared in the antibody solution and incubated at room temperature for 2 hours. After three washes, Hoechst 334422 (1:5,000, Thermo Fisher Scientific, H3570) for 10 mins, and three more 1X PBS washes. Retinas were mounted using an antifade solution (10% (v/v) mowiol (Sigma, 81381), 26% (v/v) glycerol (Sigma, G7757), 0.2M tris buffer pH 8, 2.5% (w/v) Dabco (Sigma, D27802)) on glass microscope slides with parafilm spacers to prevent tissue distortion and overlaid with coverslips (VWR, #631-0147).

### Confocal microscopy

Single cell images for deconvolution were acquired on Zeiss LSM800 microscope using a Plan-Apochromat 40X objective NA 1.4 (#420762-9900) at 1.5× digital zoom to acquire a pixel size of 0.05 µm in X and Y resolution and 0.150 µm in Z. Images were acquired from the peripheral region of the retina, in two opposing quadrants, and inner plexiform layer microglia were selected by setting the z-stack between the ganglion cell and inner nuclear layer Hoechst staining. Microglia in the ganglion cell layer were excluded due to their low population in naïve environment, and known migration from the optic nerve head to the nerve fiber layer after ONC (Heuss et al., 2018). After IPL selection, the IBA1 channel was visualized and the nearest IBA1^+^ cell was selected for imaging, thereby reducing bias in cell selection and blinded to the mitochondrial architecture within the cell. Tile scans (2×2) were necessary to acquire the area of single IPL microglia in naïve condition, resulting in images of dimension 202 x 202 µm. For each animal, 2-3 images for from two opposing quadrants were acquired. Images were stitched using Zen 2.3 or 3.5 desktop version.

Tiled (2×2) images for overview images were acquired using a Plan-Apochromat 40X objective NA 1.4 (#420762-9900) with pixel size of 0.156 µm in X and Y and 0.21 µm in Z plane for a region measuring 303 x303 µm. Tiled images (2×2) for microglia and retinal ganglion cell counting were acquired using a Plan-Apochromat 20X objective NA 0.8 (#420650-9901) with pixel resolution 0.312 x 0.312 x 0.5 µm to obtain a region measuring 605 x 605 µm in four peripheral quadrants of each retinal wholemount.

### Image processing

SVI Huygens Professional v21.10 (https://svi.nl/HomePage) was used for batch deconvolution processing of stitched confocal images. Microscope parameters for batch settings were as follows: 50 nm sampling intervals for X and Y, 150 nm in Z. The refractive index and embedding medium were set to 1.515 and 1.49, respectively. Objective quality was listed as ‘good.’ For channel settings, the back projected pinhole was set to 250 nm, and the excitation fill factor was 2. Excitation (ex) and emission (em) settings for the four channels were as follows: 405/420 nm, 488/520 nm, 561/602 nm, 633/650 nm. Deconvolution parameters were the default settings for the ‘confocal low template’ for deconvolution using the classic MLE (maximum likelihood estimation) algorithm with maximum 30 iterations.

Images were output in the Imaris file format. All confocal images presented in figures were masked in Imaris based on the IBA1-immunostained microglia of interest to remove microglia and mitochondria labeling not belonging to the analyzed cell. Supplementary videos were animated using Imaris.

### Filament tracing

Microglia morphology was traced in 3-dimensions using the *Filament* wizard in IMARIS. The starting point was detected with diameter 10 µm and 0.6 µm was used to set seeding points and the disconnected segment filtering smoothness. Final filament traces were manually edited to remove incorrect segments from the semi-automatic *Filament* tracing.

### Mitochondria and CD68 surfaces in single microglia

Surface rendering analysis was performed using IMARIS v9.1-9.3 (Bitplane, Imaris). Individual cells were cropped from the original tiled image when necessary to reduce processing time. Then, 3-dimensional surfaces were created for microglia, mitochondria and CD68 vesicles using the *Surface* wizard with surface details settings of 0.5 µm, 0.2 µm, and 0.4 µm, respectively. Each surface was manually edited to resolve any inconsistencies in surface creation. Mitochondrial surfaces were carefully edited to verify connectivity throughout the entirety of the network, often requiring manual splitting of surfaces which were joined by the thresholding within the wizard.

### Mitochondrial organelle and CD68 vesicle localization in single microglia

Completed *Surfaces* for either mitochondria or CD68 were ‘split’ using the Xtension *Split Surfaces* available via Imaris bitplane (https://imaris.oxinst.com/open/). This generates a *Split Surface* group where each individual organelle/vesicle becomes a new uniquely named *surface* nested within the *Split Surface* group. A *spot* 5 µm in diameter was manually added to identify the nucleus. Then, the *Spots and Surfaces distance* XTension calculates the distance between the nucleus *spot* and the center point of each individual *surface* in the *Split surface* group. The Xtension was modified from the original version to allow for up to 100 distance calculations and to output a *csv* file.

### Data compilation and metric calculations

Python version 3.7 with packages *pandas* (Mckinney, 2010), *matplotlib* (Hunter, 2007) and *seaborn* (Waskom, 2021) for data compilation and visualization. From Imaris, mitochondria, microglia, and CD68 surfaces, microglia filament tracings, and mitochondrial and CD68 localization *Statistics* files for each cell were exported to comma separated values file format with a unique identifier for ease in compiling cell-specific data for downstream analysis. Metric calculations for each surface type are detailed below:

#### Filament tracing

The ‘Number of Filament Sholl intersections’ from the *Statistics* file was used to plot the number of intersections per increasing radii of 1 µm (Sholl, 1953). Plotted Sholl curves were truncated at 60 µm.

#### Median mitochondrial volume and number

From the *Statistics* file, the volume of each mitochondrial *Surface* within a cell was used to determine the median mitochondrial volume per cell. Here, the total number of organelles per cell was also be extracted.

#### Sphericity

From the *Statistics* file, the sphericity metric of each mitochondrial *Surface* within a cell was used to determean the mean sphericity per cell.

#### Percentage of CD68 volume

From the *Statistics* file, the volume from each CD68 *Surface* within a cell was summed. Additionally, the sum volume of the microglia *Surface* was extracted from the microglia *Statistics* file. The percentage of CD68 volume was calculated from total CD68 volume per total microglia volume for each cell.

#### Percentage of mitochondrial volume

From the *Statistics* file, the volume from each mitochondrial *Surface* within a cell was summed. Additionally, the sum volume of the microglia *Surface* was extracted from the microglia *Statistics* file. The percentage of mitochondrial volume was calculated from total mitochondria volume per total microglia volume for each cell.

#### Mitochondrial perinuclear localization

The *Statistics* file for localization data includes the distance of each organelle from the cell soma (µm) and a unique *Surface ID* that is derived from the original mitochondrial *Surface*. From the *Statistics* file of mitochondrial volume, the *Surface ID* can be used to compile distance and volume for each unique organelle within the cell.

The percentage of mitochondrial volume in the perinuclear region was determined by calculating the sum of organelle volumes within a 10 µm radius of the cell soma, and dividing by the total mitochondria volume in that cell. The microglia soma ranges from 5-10 µm in diameter due to its non-uniform shape, therefore a 10 µm radius from the center of the soma *Spot* in IMARIS includes ∼2.5-5 µm of soma, allowing an estimation of mitochondrial localized in the perinuclear region to be at maximum 5-7 µm from the nuclear membrane.

We corroborated this perinuclear boundary distance by principal component analysis of the mitochondrial distances. Mitochondrial distances were binned in increments of 10 µm, and the sum volume in each bin was normalized to total mitochondria volume within a cell. PC loadings of the normalized values confirmed that the two bins contributing the most to principal component loadings for distances between 0-10 µm, and 71-80 µm.

### Identifying Sholl radii for PCA analysis

The Sholl curve (see Method: <u>Filament tracing)</u> is a high-dimensional, vector representation of the microglial morphology. To incorporate filament tracing into the PCA without overpowering the other scalar parameters, we first determined which Sholl radius contributed most to the differences in Sholl intersection curves. Here, Sholl intersections with 10 µm radial distances were normalized for each cell. We observed the greatest PC loadings in the 11-20 µm bin. We repeated this analysis with 5 µm radial distances and confirmed the greatest PC loadings in 11-15 µm bin. As the PC loading was greater for the 10 µm radial increments, we implemented the normalized ratio of Sholl intersections at 11-20 µm for each cell and refer to this as the Sholl index.

### Principal Component Analysis (PCA)

PCA analysis was performed using scikit-learn in Python (Pedregosa et al., 2011). Mitochondrial sphericity was omitted from analysis since it is derived from mitochondrial volume. PCA was then implemented on the remaining six metrics; median mitochondrial volume was included as a median value, while number of mitochondria, percentage of mitochondrial volume per microglia and percentage of CD68 volume per microglia volume were included as mean values for each cell. Mitochondria and CD68 localization values were included as percentage values. Data from all experimental groups were included for the PCA, where **Figure 5** shows only male and female within all conditions and genotypes and **Figure 6** also shows ovariectomized groups.

### Cell counting

RBPMS^+^-cells or cleaved-caspase3^+^/RBPMS^+^-cells were counted using the *Spots* function of Imaris v9.3, with additional manual editing. Each of these metrics was quantified from four peripheral quadrants of each analyzed retina and represented as a median value.

### Statistics

Statistical tests were performed using the Python packages *statsmodels* (Seabold and Perktold, 2010) and *Pingouin* (Vallat, 2018) as indicated in the figure legends. Groups for comparison were first tested for normal distribution and equal variance using the Shapiro test and Levene test, respectively. No data were excluded from analysis. Where applicable, post-hoc tests were performed using either Tukey’s multiple comparison of means or Conover’s test in Python packages *statsmodels* ‘multicomp’ or *Pingouin*. Significance levels are indicated using the following notation (^n.s.^ *p* > 0.05; *\*p* < 0.05; \*\**p* < 0.01; *\*\*\*p* < 0.001). For Sholl analysis comparisons, linear regression models from *lme4* package (Bates et al., 2015) were performed in R. The default contrast for unordered variables was changed to ‘contr.sum’, allowing type III ANOVA comparison. Statistical analysis and accompanying data for all figure panels are found in the **Supplementary Information.**

## Figure Legends

**Figure S1.**
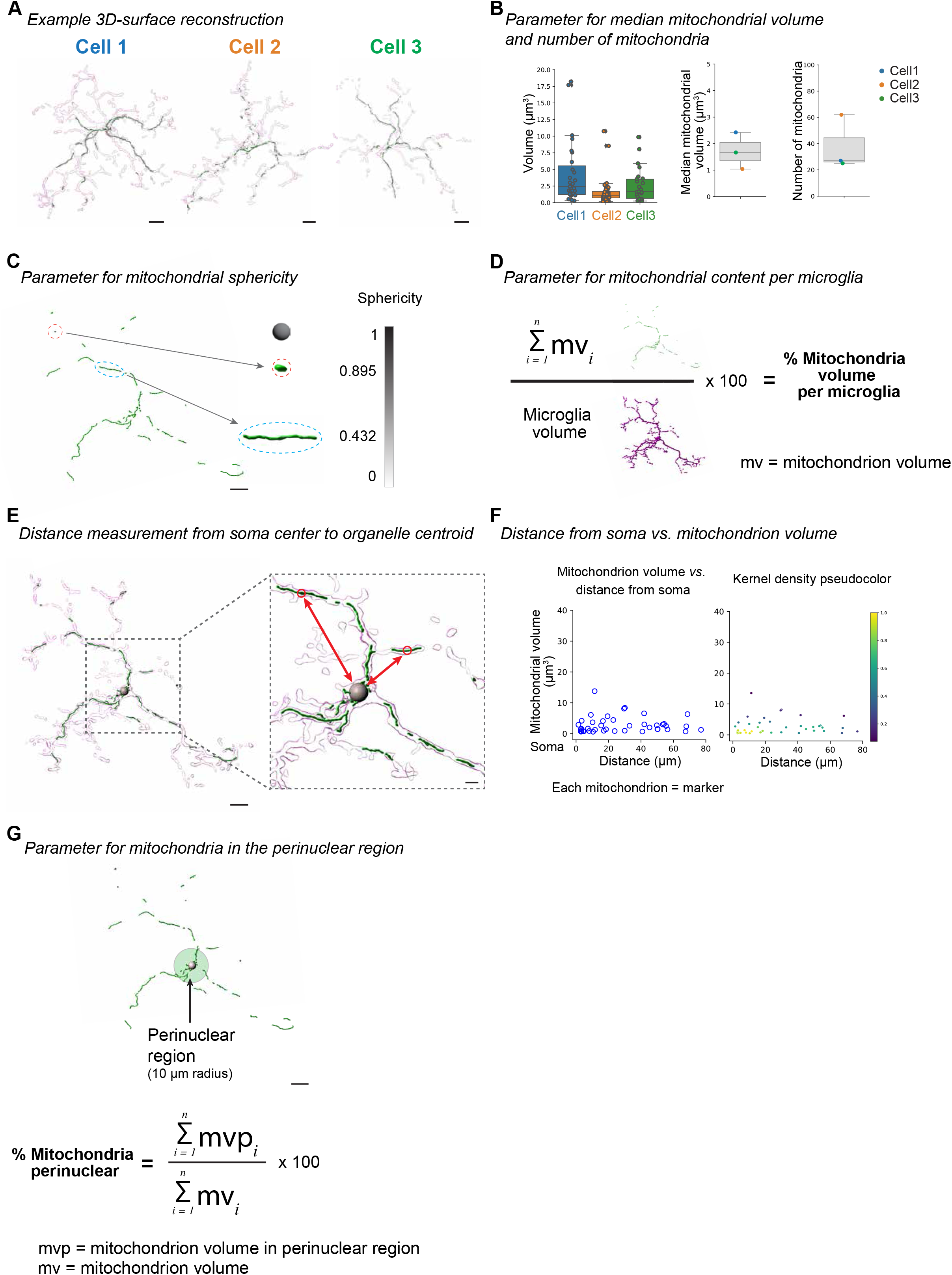
Analyzed mitochondrial parameters in microglia. Introduction to the mitochondrial parameters based on three 3D-surface reconstructions of IBA1-immunostained microglia (magenta) from IPL WT (magenta) expressing mito-Dendra2 (green, **A**). Scale bar: 10 µm. (**B**) Left, boxplot for each cell indicating the volume of every mitochondrial organelle per cell. Each dot, a mitochondrion. Center, median mitochondrial volume and right, number of mitochondria for each cell at left. Boxplot minimum and maximum: InterQuartile Range (IQR) around median (center line). Whiskers: 1.5 IQRs. Black diamond points: outliers outside 1.5 IQRs. (**C**) Mitochondrial sphericity. Left, example mitochondrial 3D-surface (green) from a WT microglia. Ratio of mitochondrion volume to mitochondrion surface area represents sphericity with two exemplar organelles (orange, sphericity =0.895) and elongated (blue, sphericity = 0.432). Sphericity is represented as a mean sphericity value of all mitochondria per cell. Scale bar: 15 µm. (**D**) Percentage of the sum of mitochondrial volume in a cell (**Figure S1B, left**) over the microglial volume represents mitochondrial content per cell. (**E**) Vesicle distance measurement. Example 3D-volume surfaces of an IBA1-immunostained microglia (magenta) with mito-Dendra2 expression (green). Grey sphere: cell soma. Right, zoom-in of region of interest (dashed line). Example red circles: centroid of mitochondrion. Arrows: distance from centroid to soma determined from the centroid of each mitochondrion/vesicle to the center of the soma-sphere (red arrows). Scale bar: 15 µm, zoom-in: 5 µm. (**F**) Mitochondria or CD68 vesicle localization parameter exemplified using mitochondrial localization. Scatterplot depicts the distance from the soma (µm) *vs.* the mitochondrial volume (µm^3^) for each mitochondrion shown in (**Figure S1E**). To illustrate overlapping mitochondria, scatterplot markers were pseudo-colored with kernel density estimations. (**G**) Example mitochondrial surface (green) from microglia depicted in (**Figure S1E**). The perinuclear region (shaded green sphere) was determined as the region within a 10 µm radius from the cell soma point. The percentage of mitochondria in the perinuclear region is the sum volume within the perinuclear region over the total mitochondrial volume. Scale bar: 15 µm.

**Figure S2.**
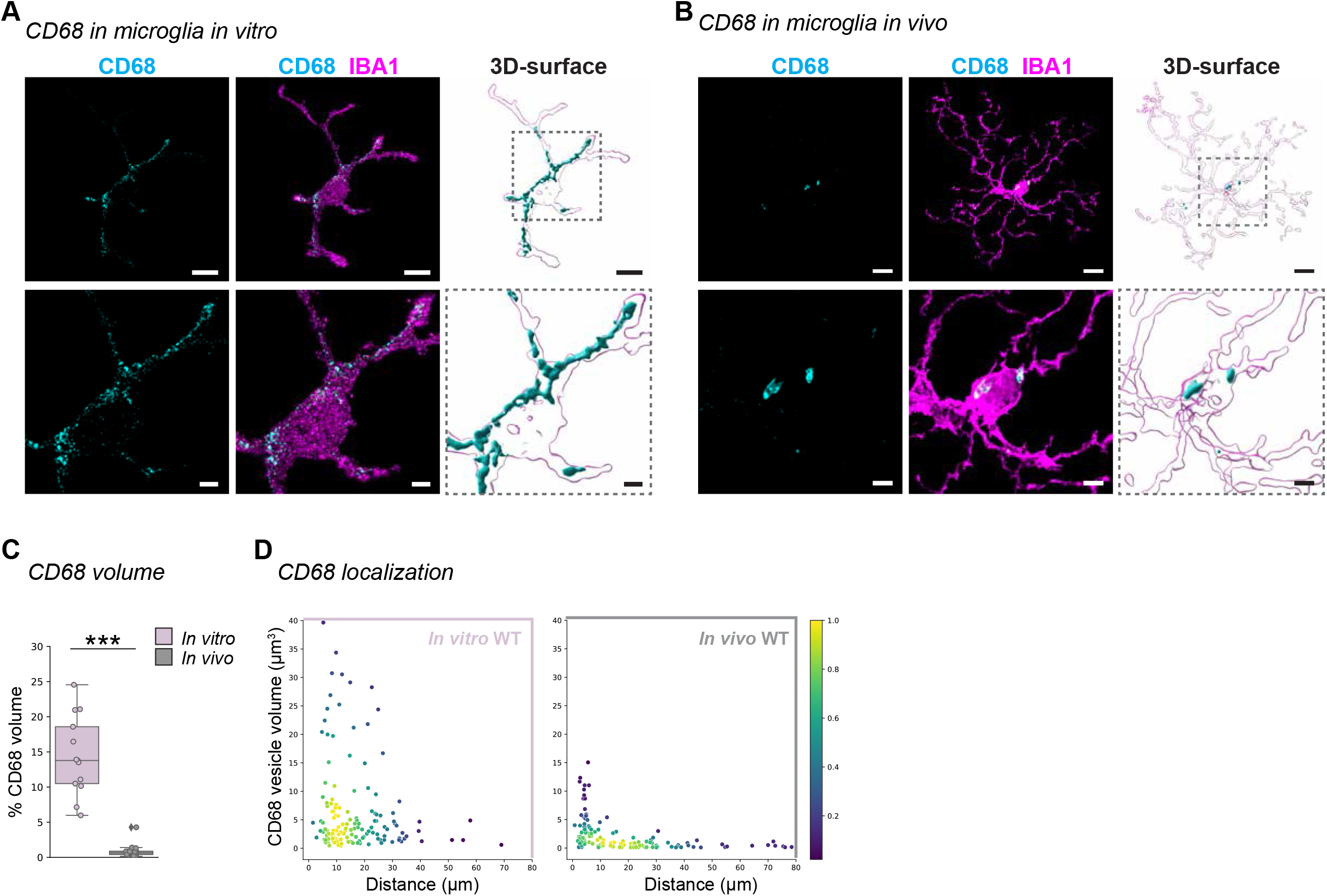
Microglia *in vitro* exhibit increased CD68 expression. (**A**-**B**) IBA1-immunostained microglia (magenta) from Figure 1F-G co-labeled with CD68 (cyan) from *in vitro* (**A**) or *in vivo* microglia (**B**). Next, corresponding 3D-surfaces. Below: Zoom-in of region of interest (dashed outline) from image and 3D-surface. Scale bar: 10 µm, zoom-in: 3 µm. (**C**) Percentage of total CD68 volume per microglia volume. Boxplot minimum and maximum: InterQuartile Range (IQR) around median (center line). Whiskers: 1.5 IQRs. Black diamond points: outliers outside 1.5 IQRs. Wilcoxon rank sum test: *p* < 0.0001. (**D**) CD68 vesicle localization. Scatterplot depicting CD68 vesicle volume *vs.* distance from the cell soma (0, origin) of the population of vesicles in *in vitro* (left, dusty pink) and *in vivo* (right, grey) conditions. Point density, pseudo-colored. *** p < 0.001. See **Supplementary Information** for experimental, retina and cell numbers, statistical tests and corresponding data.

**Figure S3.**
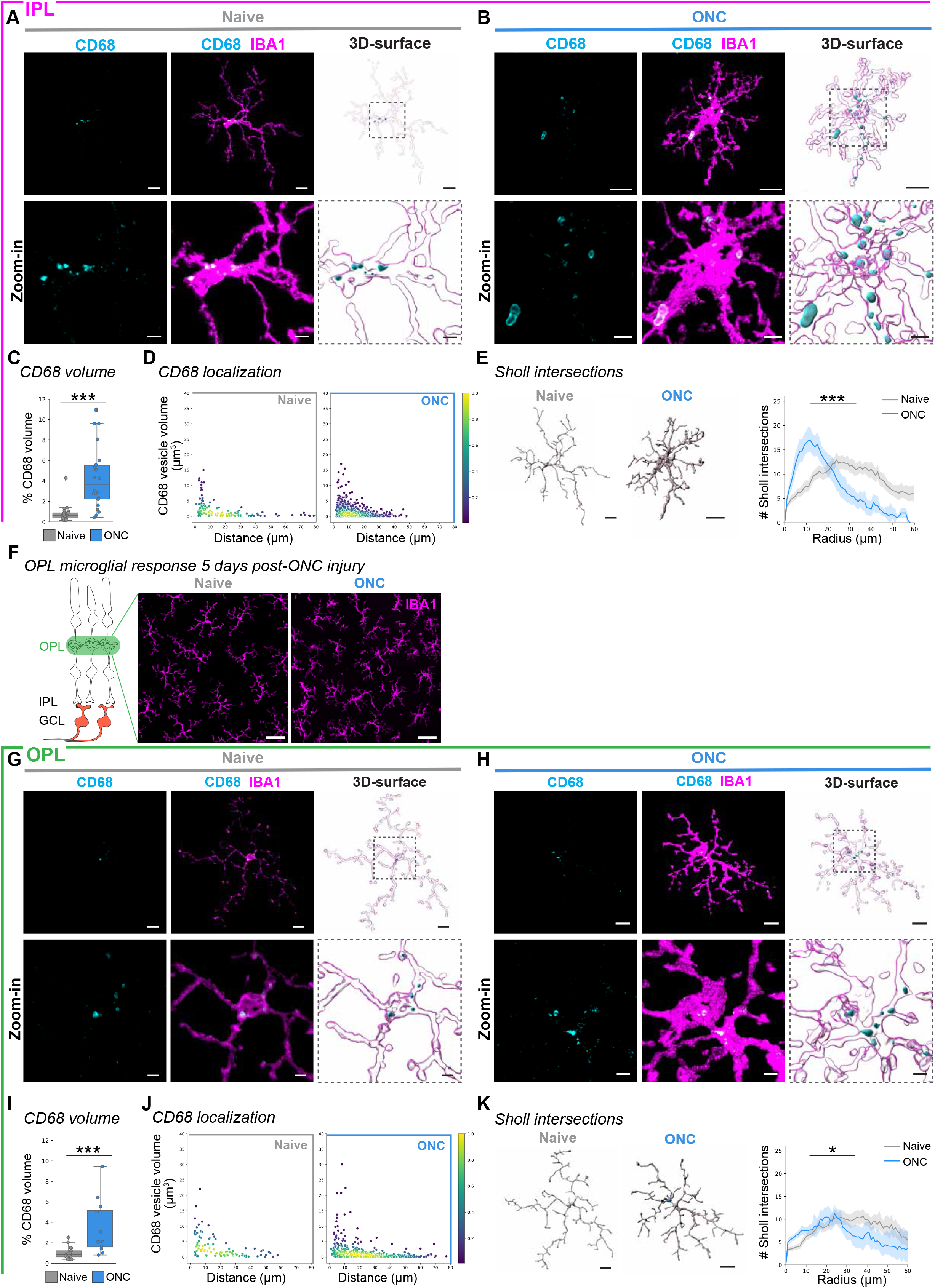
CD68 and microglial morphology changes in ONC-responsive IPL and OPL microglia (**A-E**) Microglial response in IPL WT microglia. (**A**-**B**) Representative IBA1-immunostained IPL microglia (magenta) from Figure 3D-E co-labeled with CD68 (cyan) in naïve WT (**A**, grey) or ONC WT microglia (**B**, blue). Next, corresponding 3D-surfaces. Below: Zoom-in of region of interest (dashed line) from image and 3D-surface. Scale bar: 10 µm, zoom-in: 3 µm. (**C**) Percentage of total CD68 volume per microglia volume. Boxplot minimum and maximum: InterQuartile Range (IQR) around median (center line). Whiskers: 1.5 IQRs. Black diamond points: outliers outside 1.5 IQRs. Wilcoxon rank sum test: *p* < 0.0001. (**D**) CD68 vesicle localization. Scatterplot depicting CD68 vesicle volume *vs.* distance from the cell soma (0, origin) of the population of vesicles in naïve WT (left) and ONC WT (right) conditions. Point density, pseudo-colored. (**E**) 3D-filament tracings of microglia from naïve WT (left) or ONC WT (right). Scale bar: 10 µm. Line plot for mean number of Sholl intersections per radial distant from the soma (µm) with 95% confidence interval band. Linear mixed effects model: *p* < 0.0001. (**F**) Left, retinal side-view schematic. OPL, outer plexiform layer. IPL, inner plexiform layer. GCL, ganglion cell layer. Overview confocal image of immunostained OPL microglia (IBA1, magenta) in naïve WT (left) and 5 days after ONC WT (right) condition. Scale bar: 30 µm. (**G**-**K**) Microglial response in OPL WT microglia. (**G**-**H**) CD68 (cyan) immunostaining in the IBA1-immunostained OPL microglia (magenta) from Figure 2I-**J** for naïve WT microglia (**G**) or ONC WT microglia (**H**). Next, corresponding 3D-surfaces. Below: Zoom-in of region of interest (dashed outline) from image and 3D-surface. Scale bar: 10 µm, zoom-in: 3 µm. (**I**) Percentage of total CD68 volume per microglia volume. Boxplot minimum and maximum: InterQuartile Range (IQR) around median (center line). Whiskers: 1.5 IQRs. Black diamond points: outliers outside 1.5 IQRs. Wilcoxon rank sum test: *p* = 0.0007. (**J**) CD68 vesicle localization. Scatterplot depicting CD68 vesicle volume *vs.* distance from the cell soma (0, origin) of the population of vesicles in naïve (left, grey) and ONC (right, blue) conditions. Point density, pseudo-colored. (**K**) 3D-filament tracings of microglia from naïve WT (left) or ONC WT (right). Scale bar: 10 µm. Line plot for mean number of Sholl intersections per radial distant from the soma (µm) with 95% confidence interval band. Linear mixed effects model: *p* = 0.0302. * p < 0.05, *** p < 0.001. See **Supplementary Information** for retina and cell numbers, statistical tests and corresponding data.

**Figure S4.**
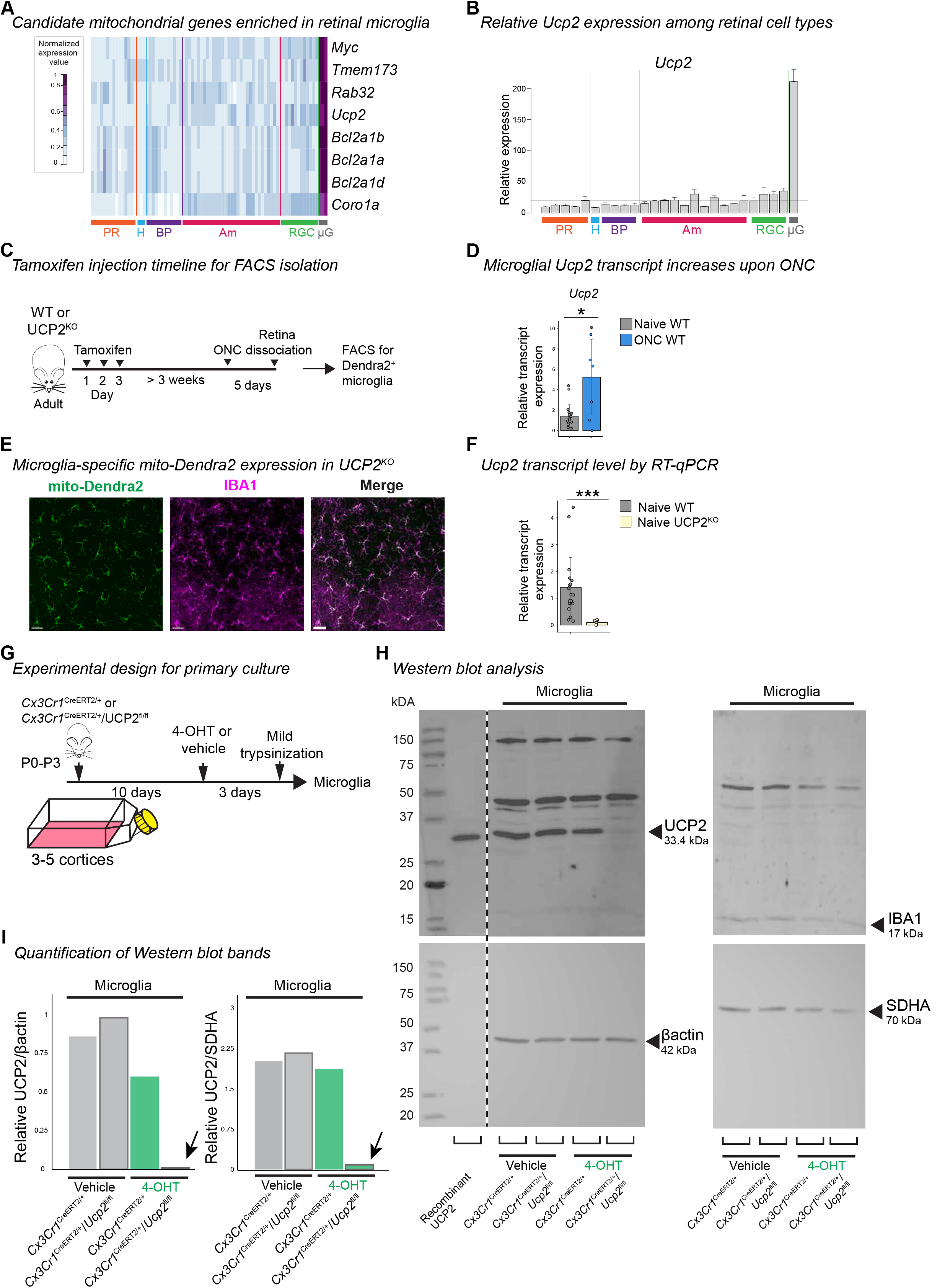
Selective expression and knockout of UCP2 in retinal microglia. (**A**-**B**) Relative fold change expression level of mitochondrial-associated genes (**A**) and *Ucp2* (**B**) among retinal cell types. Plotted from available data in *Siegert et. al*. (1). Purple and blue: high and low normalized expression value, respectively. PR: photoreceptors, H: horizontal cells, BP: bipolar cells, Am: amacrine cells, RGC: retinal ganglion cells, µG: microglia. (**C**) Timeline for tamoxifen administration to induce mito-Dendra2 expression or in combination UCP2^KO^ in microglia used for FACS (fluorescence activated cell sorting). (**D**) Boxplot of relative *Ucp2* transcript expression from Dendra2^+^-FACSed retinal microglia in naïve WT or 5 days ONC WT. Boxplot minimum and maximum: InterQuartile Range (IQR) around median (center line). Whiskers: 1.5 IQRs. Black diamond: outliers outside of 1.5 IQRs. Naïve: grey. ONC: blue. Wilcoxon rank sum test: *p* = 0.0474. (**E**) Microglial specificity of mito-Dendra2 labeling in UCP2^KO^ mouse model. Overview image of immunostained microglia (IBA1, magenta) in the inner plexiform layer (IPL) expressing the mitochondrial label (Dendra2, green) in retinal wholemounts. Scale bar: 50 µm. (**F**) Relative *Ucp2* transcript expression from tamoxifen-induced naïve WT or naïve UCP2^KO^ FACSed-microglia from the retina. Wilcoxon rank sum test: *p* < 0.0001. (**G**) Experimental timeline for primary murine mixed glial cultures to obtain microglia and astrocyte-enriched cultures for protein analysis. P0-P3: postnatal day 0 to 3. 4-OHT: 4-hydroxytamoxifen. Next: IBA1-immunostained (magenta) mixed glial culture at day 10 prior to trypsinization. Scale bar: 50 µm. (**H**) Western blot analysis of UCP2 protein expression (left, top), β-actin loading control (left, bottom), IBA1 microglial marker (right, top), or SDHA (Succinate dehydrogenase complex flavoprotein subunit A) mitochondrial loading control (right, bottom) in primary microglia of either *Cx3cr1*^CreERT2/+^ or *Cx3cr1*^CreERT2/+^/UCP2^fl/fl^ with vehicle or 4-OHT (n = 1). kDa, kilodaltons. (**I**) Quantification of UCP2 band intensity relative to β-actin loading control (left) or relative to SDHA (right) mitochondrial loading control. * p < 0.05, *** p < 0.001. See **Supplementary Information** for retina and cell numbers, statistical tests and corresponding data.

**Figure S5.**
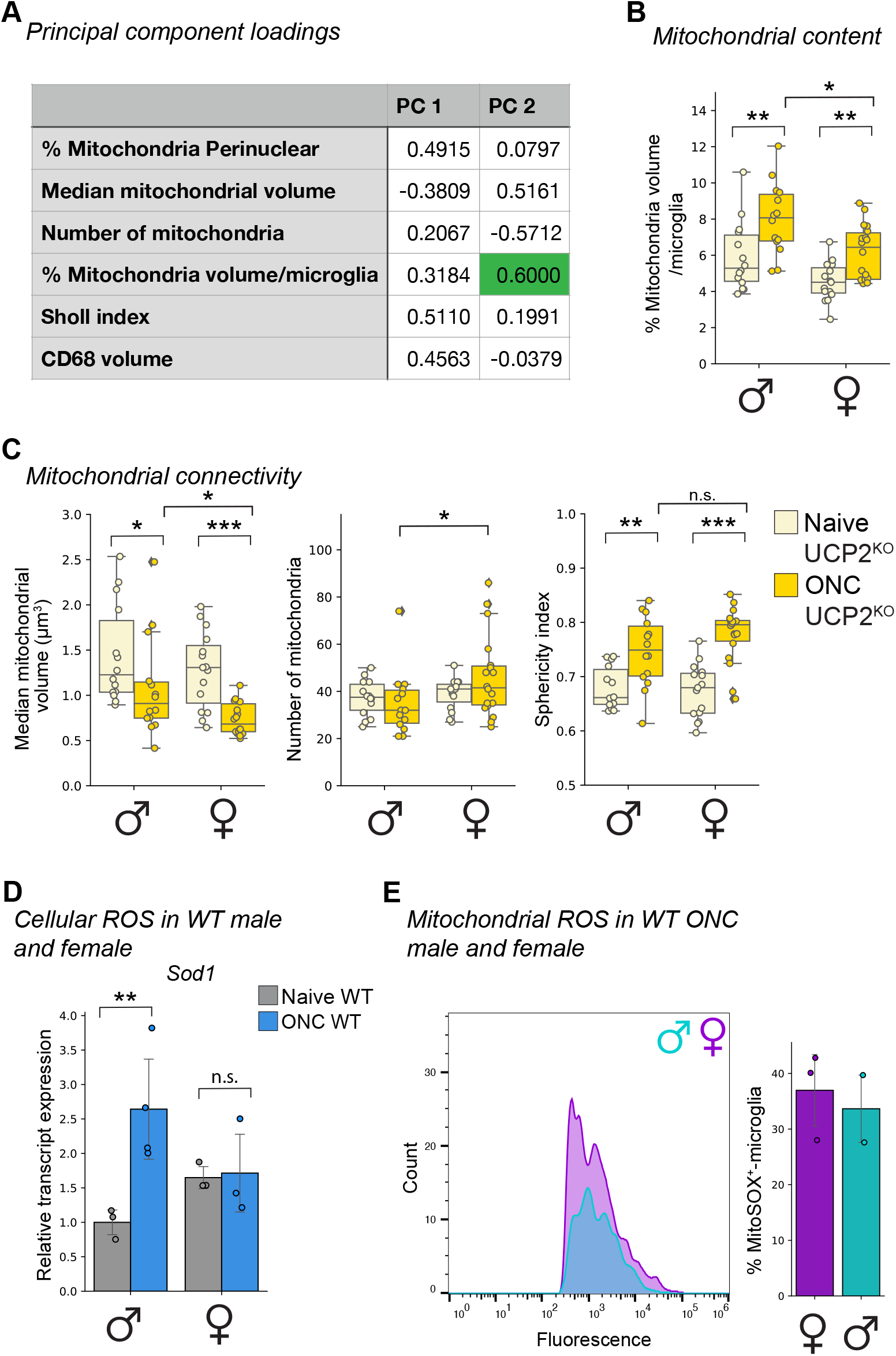
Sex-separated UCP2^KO^ mitochondrial parameters. (**A**) Table of principal component (PC) loadings from Figure 5. Green highlight: Highest loading for PC2. (**B**-**C**) Mitochondrial parameters. Boxplot minimum and maximum: InterQuartile Range (IQR) around median (center line). Whiskers: 1.5 IQRs. Black diamond: outliers outside 1.5 IQRs. Each overlaid point represents data point of a single microglia. Naïve UCP2^KO^: tan. ONC UCP2^KO^: gold. (**B**) Percentage of mitochondrial volume per microglial volume. Kruskal-Wallis test: *p* < 0.0001. Selected Conover’s post-hoc comparisons with Holm p-adjustment: Naïve ♂UCP2^KO^ *vs.* ONC ♂UCP2^KO^, *p* = 0.0076; Naïve ♀UCP2^KO^ *vs.* ONC ♀UCP2^KO^, *p* = 0.0073; ONC ♂UCP2^KO^ *vs.* ONC ♀UCP2^KO^, *p* = 0.0266. (**C**) Mitochondrial network connectivity determined by median mitochondrial volume (left, Kruskal-Wallis test: *p* < 0.0001. Selected Conover’s post-hoc comparisons with Holm p-adjustment: Naïve ♂UCP2^KO^ *vs*. ONC ♂UCP2^KO^, *p* = 0.0348; Naïve ♀UCP2^KO^ *vs.* ONC ♀UCP2^KO^, *p* < 0.0001; ONC ♂UCP2^KO^ *vs.* ONC ♀UCP2^KO^, *p* = 0.0303), number of organelles (center, Wilcoxon rank sum test: *p* = 0.0383) and mean sphericity (right, Kruskal-Wallis test: *p* < 0.0001. Selected Conover’s post-hoc comparisons with Holm p-adjustment: Naïve ♂UCP2^KO^ *vs.* ONC ♂UCP2^KO^, *p* = 0.0029; Naïve ♀UCP2^KO^ *vs.* ONC ♀UCP2^KO^, *p* < 0.0001; ONC ♂UCP2^KO^ *vs.* ONC ♀UCP2^KO^, *p* = 0.2323). (**D**) Bar plot depicting relative *Sod1* transcript expression from FACSed mito-Dendra2^+^ microglia in naïve WT and ONC WT male and female microglia. Transcript expression relative to male naïve WT. Kruskal-Wallis test: *p* = 0.0269. Selected Conover’s post-hoc comparisons with Holm p-adjustment: ♂WT Naïve *vs*. ♂WT ONC: *p* = 0.0026, ♀WT Naïve *vs.* ♀WT ONC: *p* = 0.8554. (**E**) Frequency plot of MitoSOX fluorescence from FACSed mito-Dendra2^+^ retinal microglia for WT ONC male (turquoise) and female (purple). Corresponding bar plot of percentage of microglia that are MitoSOX^+^. ** p < 0.01, *** p < 0.001, ^n.s.^ p > 0.05: not significant. See **Supplementary Information** for retina and cell numbers, statistical tests, other post-hoc comparisons and corresponding data.

**Table S1.**
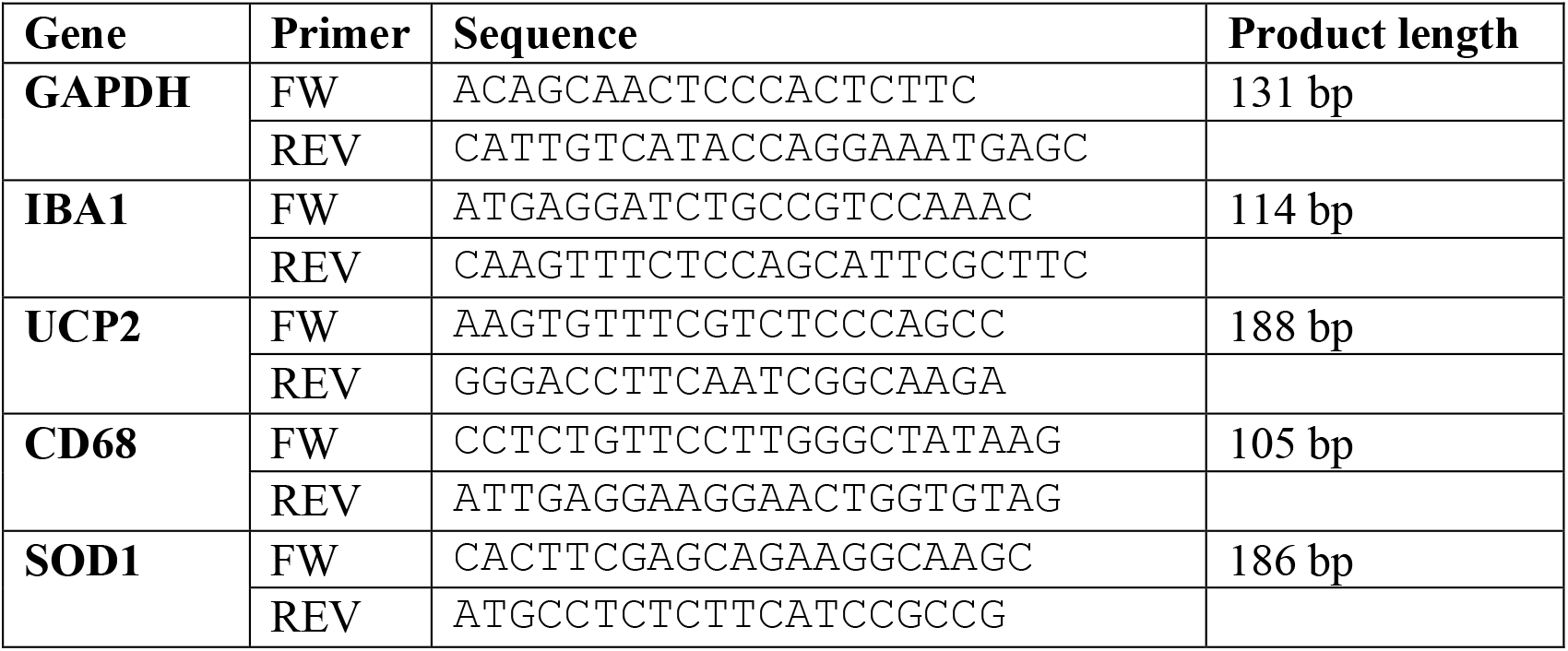
Primer sequences.

**Table S2.** Antibody table.

| Antibody | Dilution | Source | Catalog No. | RRID |
| --- | --- | --- | --- | --- |
| Rabbit anti-IBA1 | 1:500 | GeneTex | GTX100042 | AB_1240434 |
| Goat anti-IBA1 | IHC: 1:250<br>WB: 1µg/ml | Abcam | ab 5076 | AB_2224402 |
| Rat anti-CD68 | 1:500 | Bio-Rad | MCA1957 | AB_322219 |
| Guinea pig anti-RBPMS | 1:200 | PhosphoSolutions | 1832-RBPMS | AB_2492226 |
| Rabbit anti-RBPMS | 1:200 | Abcam | ab194213 | AB_2920590 |
| Rabbit anti- Cleaved caspase 3 | 1:400 | Cell Signaling Technology | #9661S | AB_2341188 |
| UCP2 | 1:1000 | Pohl Lab, purified at Pineda | NT-Tier2 | n/a |
| SDHA | 1:5000 | Abcam | ab14715 | AB_301433 |
| β-actin | 1:5000 | Sigma | A5441 | AB_476744 |
| Goat anti-rabbit IgG (H+L) Alexa Fluor 568 | 1:2000 | ThermoFisher | A11036 | AB_10563566 |
| Goat anti- rat IgG (H+L) Alexa Fluor 647 | 1:2000 | ThermoFisher | A21247 | AB_141778 |
| Donkey Anti-Guinea Pig IgG (H+L) Alexa Fluor 647 | 1:1000 | Jackson ImmunoResearch | 706-605-148 | AB_2340476 |
| Donkey anti-rabbit IgG (H+L) Alexa Fluor 568 | 1:2000 | ThermoFisher | A10042 | AB_2534017 |
| Donkey anti-goat IgG (H+L) | 1:2000 | ThermoFisher | A11057 | AB_2534104 |
| Donkey anti-goat Alexa Fluor 568 IgG (H+L) Alexa Fluor 488 | 1:2000 | ThermoFisher | A11055 | AB_2534102 |
| Donkey anti-rabbit IgG (H+L) Alexa Fluor 488 | 1:2000 | ThermoFisher | A21206 | AB_2535792 |
| Donkey anti-Rat IgG (H+L) Antibody DyLight 650 | 1:2000 | ThermoFisher | SA5-10029 | AB_2556609 |
| Donkey anti-Rat IgG (H+L) Alexa Fluor 488 | 1:2000 | ThermoFisher | A21208 | AB_2535794 |
| Donkey anti-rabbit IgG (H+L) Alexa Fluor 647 | 1:2000 | ThermoFisher | A31573 | AB_2536183 |

**Supplementary Information**. Statistical analysis.

**Supplementary Video 1**. 3D-surface reconstruction of in vitro microglia and mitochondria from **Figure 1F**.

**Supplementary Video 2**. 3D-surface reconstruction of in vivo microglia and mitochondria from **Figure 1G**.

**Supplementary Video 3**. 3D-surface reconstruction of male UCP2KO microglia and hyperfused mitochondria in **Figure 5D**.

